# DNA Divers: volunteer-based eDNA capture for local and global marine biodiversity monitoring

**DOI:** 10.1101/2025.05.26.656130

**Authors:** Erika F. Neave, Annie Watson, Alice Cunnington, Giulia Maiello, Natasha Yates, Candice J. Parkes, Fiona Crouch, Harry J. N. Catherall, Peter Shum, Wang Cai, Romain Allemann, Karen Boswarva, Rosie Horner, Wendy Northway, Frederic Bezombes, Charlotte Bolton, Bob Anderson, Ela Johnson, Sarah Meek, Grant Smith, Stefano Mariani

## Abstract

Volunteer-based biodiversity recording is a powerful source of scalable data yet to be used to its full potential by the scientific community. Coastal ecosystems are varied and diverse, making it difficult for managers to identify flexible methods for monitoring biological components. Emerging technologies such as environmental DNA (eDNA) analysis are promising for measuring aquatic biodiversity, yet most management organizations lack personnel and capacity to collect molecular data.

Here we investigate, together as professional and non-professional scientists, the efficacy of a quasi-passive eDNA capture technique. Volunteer SCUBA divers and snorkelers used low-cost materials, namely cotton medical gauze, to capture eDNA via swimming. We compared this method to conventional eDNA capture techniques in an aquarium and nature, collectively iterating the field and laboratory procedures to improve feasibility.

With a small (< 30) network of volunteers, we detected 275 unique teleost and elasmobranch species from varied marine habitats spanning 90 degrees of latitude. The swimming motion of divers was more effective than stationary soaking and the fish communities sampled by divers were comparable to that of conventional eDNA samples.

The ease of this technique matched by the eagerness and generosity of volunteers presents an untapped, viable approach for scaling multi-species marine eDNA monitoring as well as an avenue for improving science literacy.

**Data availability:** https://zenodo.org/records/14853820?token=eyJhbGciOiJIUzUxMiJ9.eyJpZCI6ImYxMTlhODFiLTI3NGYtNDQ2Ni05MDYyLTIyOTlkNWVhY2FkNiIsImRhdGEiOnt9LCJyYW5kb20iOiIwYmZlMzJhMWU0YWZkM2ZhMWEwYTE0NmY0ZWM5YWZkNSJ9.0R7GJjVJLEuq-UJVKA1CcMT8LCTamv3GEvPl-5ItzP10pFRd-cAzAr_Q5ggjEewZUtByHjBu-eSegWWlLAx_Rw

## Introduction

Marine and coastal habitats modulate processes affecting water quality, disease, and climate, whilst providing many ecosystem services that bolster the economy, such as fisheries, tourism, and recreation ^1^. Biodiversity loss causes the health of these habitats to diminish and ecosystem services to wane ^2,3^. Marine environmental DNA (eDNA) analyses have the accuracy and sensitivity necessary to monitor a variety of species across the tree of life (i.e. microbes to macro-organisms) ^4^. Yet, despite substantial advances in methodology, one major roadblock for the uptake of eDNA monitoring is the ability to capture eDNA efficiently and at scale. Particularly in marine environments, eDNA exists in trace amounts such that water filtration, even when assisted mechanically by vacuum pumps or autonomous samplers ^5^, can require considerable time investment and bespoke, costly instruments ^6,7^.

Despite the challenge of scalability, eDNA analysis is a promising tool for marine management and conservation because in many cases it offers advantages over conventional methods. For instance, morphological identification methods (e.g., visual surveys, video, plankton tow-net) require expert taxonomists, visible conditions and exclude organisms that cannot be seen with the naked eye or distinguished using microscopy. Moreover, physical survey methods (e.g., trawling, electrofishing), in addition to visual identification constraints, can damage delicate habitats and cause disturbance or require the collection of specimens, which can be counterproductive in conservation contexts. In marine environments, eDNA analysis often results in higher species richness compared to visual dive surveys and can even differentiate communities in habitats over short distances (< 100 m) ^8,9^. When comparing tandem trawl data to eDNA, both methods result in similar species richness and the relative abundance of eDNA sequence reads correlate with trawl catch counts ^10,11^. The scientific community continues to demonstrate the advantages of eDNA analyses compared to conventional monitoring methods which is important for understanding how eDNA data can enhance and be integrated with traditional measurements ^12,13^. Nonetheless, mounting evidence supports the adoption of eDNA analysis for holistic marine biodiversity monitoring, but strategies to do so have yet to be formalized and implemented ^14^.

One of the largest and fastest growing sources of biodiversity data comes from the grass-roots efforts of volunteer monitoring programs or citizen science ^15^. Citizen science, or the inclusion of non-professional scientists in the scientific process, is a methodology employed across a variety of fields to enhance the reach and feasibility of research by covering areas that would otherwise be too extensive for traditional research projects alone. For example, crowdsourcing, particularly through recruitment on web platforms (i.e., Zooniverse, SciStarter, iNaturalist), encourages volunteers to collect and analyze large volumes of data. This concept has been used to scale up eDNA monitoring, for instance water samples from the entire coastline of Denmark were collected by volunteers at coordinated time points, providing snapshots of coastal fish biodiversity^16^. Similarly, eDNA citizen science projects have emerged in an effort to cover large geographic areas and establish long-term monitoring strategies, such as the California environmental DNA (CALeDNA) program ^17^ and UNESCO-led expeditions of marine world heritage sites.

Since the process of actively filtering eDNA requires time and investment in trained personnel, passive eDNA capture techniques have been explored as an alternative ^18–20^. The first study on passive eDNA capture tested the ability of porous materials (i.e., montmorillonite clay and granular activated carbon) to adsorb extracellular DNA and found that after soaking for days to weeks the materials accrued a similar amount of DNA to 1 L of filtered water ^19^. In nature, marine eDNA was captured on filter membranes (i.e., suspended in the water column for up to 24 hours) and compared to actively filtered seawater; this study resulted in the detection of 172 fish taxa, 109 of which were detected passively on the suspended membranes ^18^. Despite these passive eDNA capture techniques requiring deployment over hours to days the apparatus could be left unattended, meaning that the researchers spent their time on other tasks and that there was limitless potential for replication, contrary to filtering eDNA from water which requires more time investment with increased replication. To date most passive eDNA sampling studies have focused on controlled conditions ^20,21^, yet preliminary studies in nature show promise for large-scale sampling ^18,22^.

While studies demonstrate that citizen science and passive eDNA sampling can help scale marine eDNA projects ^16,18^, these solutions have only been applied independent of each other. One promising branch of eDNA capture research has involved combining passive sampling, using sterile cotton in a perforated 3D-printed spherical probe (the ‘metaprobe’), with conventional trawl surveys by placing the metaprobe into the fishing net, detections from which resembled species present in the catch or native to the area ^22,23^. Here we combine the metaprobe passive eDNA sampler with the power of SCUBA diving citizen scientists to scale up biodiversity monitoring efforts in coastal marine habitats (Fig. 1). First, we evaluate the influence of metaprobe soaking time in an aquarium; then we assess the influence of different preservation and DNA extraction techniques, testing the approach in natural marine habitats, and ultimately comparing the results to conventional water filtration (Fig. 1). Finally, through engagement with existing citizen science networks, independent volunteers, SCUBA diving clubs, the dive charter industry, non-profits, and statutory governing agencies, we scale up, sampling at dive sites spanning four continents (Fig. 1, table 1).

**Fig 1.**
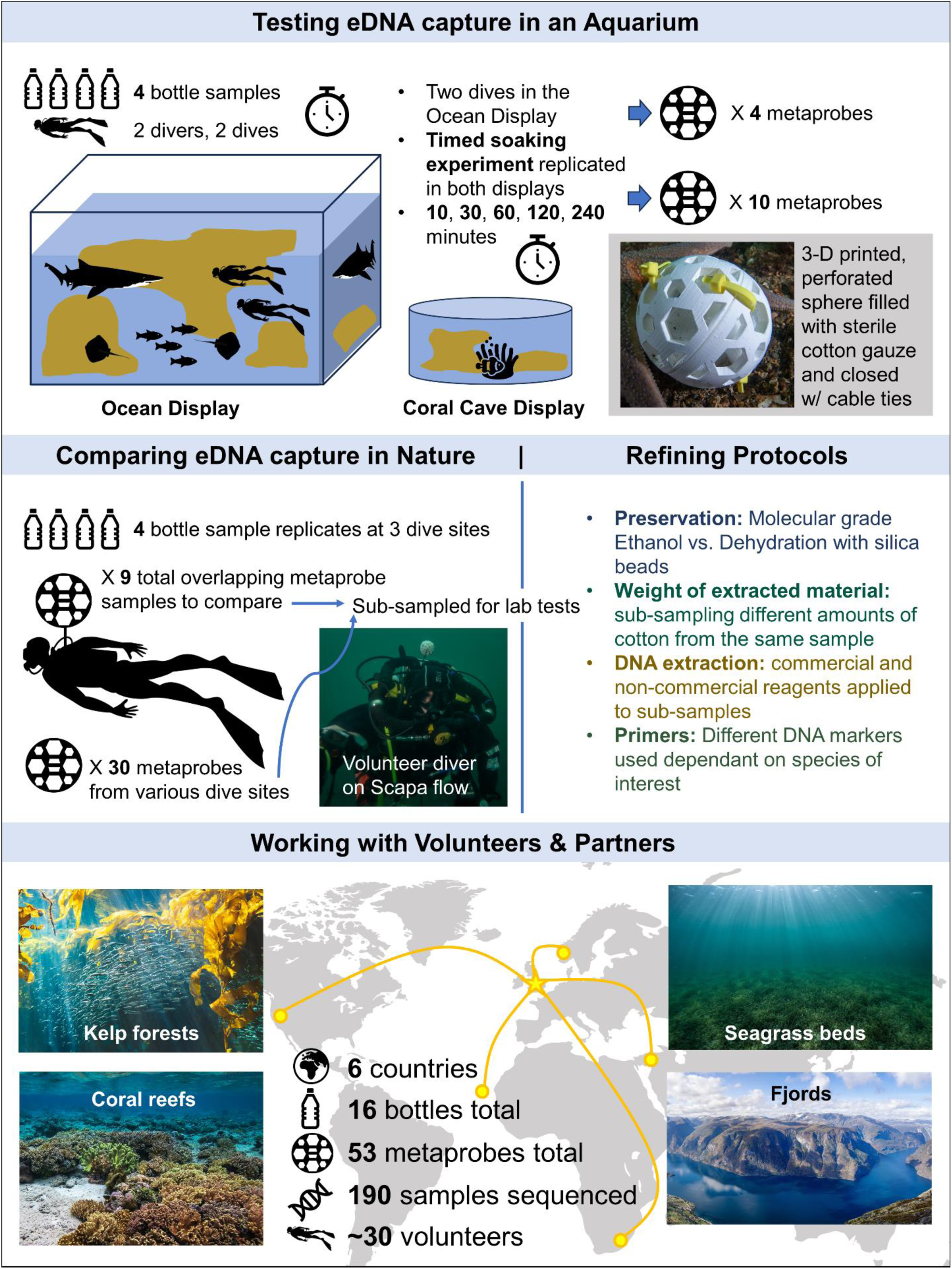
Schematic of the DNA divers citizen science project.

**Table 1.**
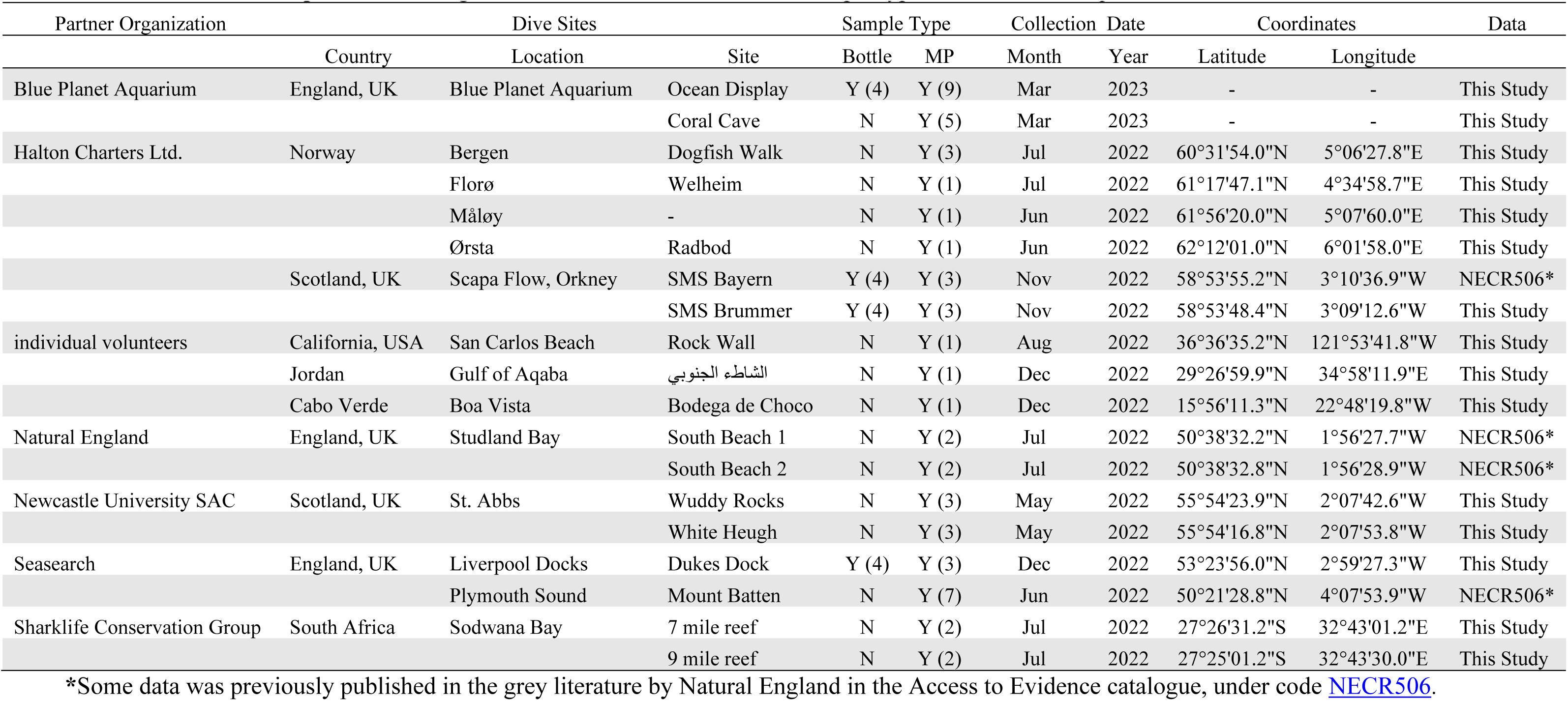
Dive sites sampled. Partner organizations, dive site location, and sample type information and quantities.

## Methods and Materials

### Sample Collection

Sample collection events took place at various dive sites in the UK, Cape Verde, Jordan, Norway, South Africa, and the USA as well as the Blue Planet Aquarium (Table 1). In some locations eDNA sampling by water filtration was performed to compare to metaprobes which had been worn by snorkelers or SCUBA divers. Metaprobes were 3-D printed using polylactic acid (PLA) and where possible were prepared in the laboratory and distributed to snorkelers and SCUBA divers. Metaprobes were filled with between one to three rolls of sterile cotton medical gauze, and through practice we established that two whole rolls of medium-sized cotton gauze rolls (10 cm x 3.7 m) were sufficient for sub-sampling. Water samples were collected in bleach-cleaned 1.5 L bottles and were filtered on site through a 0.45 µm Sterivex filter (PES membrane, Merck Millipore). In the Blue Planet Aquarium, both sample types were collected (Table 1) alongside a controlled timed experiment, where metaprobes were soaked in display tanks (Fig. 1). The timed experiment was repeated in two independent tanks (the Ocean exhibit and the Coral Cave). In each tank, five metaprobes were placed in a mesh dive bag and allowed to soak at the surface of the tanks; one was removed and preserved at each time point: 10, 30, 60, 120, and 240 minutes. Sterivex filters were dried by pushing air through the cartridge. Metaprobe gauze was either dried using silica beads or stored in 100% molecular grade ethanol. All samples were transported in an insulated box with ice packs until they could be stored at -20°C. Detailed metaprobe preparation instructions are in section A of the supplementary methods.

### DNA extraction and sequencing

#### DNA extraction

All combinations of DNA extraction and preservation techniques, which resulted in 190 sub-samples not including controls, can be found in Table S1. DNA was extracted from the metaprobe gauze and sterivex filters using either the Qiagen DNeasy Blood and Tissue kit following the manufacturer’s protocol, but with slight modifications (see Supplementary Text), or ‘Mu-DNA’, a non-commercial published modular protocol, from which we combined steps from the ‘water’ and ‘tissue’ variants (see section B of the supplementary methods) ^29^. To remove the sterivex filters from the capsules the plastic was broken open using pliers and the center column holding the filter was emptied onto a petri dish. The filter was then removed from the center column of the capsule using tweezers. Half of the filter was saved as archival material, and the other half was cut into small pieces using dissecting scissors and placed into a 1.5 ml Eppendorf tube. The metaprobe cotton gauze was prepared by cutting off small pieces from various parts of the larger roll. The small pieces were placed onto blotting paper to remove ethanol and were weighed in weighing boats. The various weight ranges were established based on the amount of cotton that could reasonably fit in different sized Eppendorf tubes: heavy (0.9 - 1.1 g), medium (0.6 – 0.8 g), and standard (0.2 – 0.45 g). Each weight range was treated with a proportional volume of lysis buffer (i.e., 5000 µl, 3000 µl, 1000 µl). All non-disposable equipment used for DNA extractions was soaked in a 10% bleach solution, followed by 5% Lipsol detergent then Milli-Q water, and UV treated for at least 30 minutes. Disposable equipment used for DNA extractions was sterile and exposed to ultra-violet light under a hood for 30 minutes before use.

#### DNA amplification

Three replicates of each extract were amplified by polymerase chain reaction (PCR). The tele02 (specifically: forward sequence Tele02-F 5’-AAACTCGTGCCAGCCACC-3’ and the reverse sequence Tele02-R 3’-GGGTATCTAATCCCAGTTTG-5’) or elas02 (specifically: forward sequence Elas02-F 5’-GTTGGTHAATCTCGTGCCAGC-3’ and the reverse sequence Tele02-R 3’-CATAGTAGGGTATCTAATCCTAGTTTG-5’) primers were used to target 167 and 171 bp regions of the 12S gene, depending on the sample library ^36^ (Table S1). The PCR cycles for tele02 were as follows: 95°C for 10 min, followed by 35 cycles of 95°C for 30 s, 60°C for 45 s, 72°C for 30 s, and finishing at 72°C for 5 min followed by a 4°C hold. The PCR cycles for elas02 were as follows: 94°C for 5 min, followed by 35 cycles of 94°C for 1 min, 54°C for 1 min, 72°C for 1 min, and finishing at 72°C for 5 min followed by a 4°C hold. From 16 syringe-filtered samples and 53 metaprobes (sub-sampled for protocol refinement), 190 sub-samples were sequenced not including controls such as field blanks (aquarium N=3, Orkney N=6, Liverpool N=3), extraction blanks (per extraction batch N=12), PCR blanks (all sequencing runs N = 12) and positive controls (all sequencing runs N = 12). All PCR replicates were visualized on a 2% agarose gel (150 ml 1X TBE buffer with 3 g agarose powder) stained with 1.5 μl SYBRsafe dye (InVitrogen). The PCR replicates were then combined by sample and purified using 1:1 ratio of PCR product to Mag-Bind® Total Pure NGS magnetic beads (Omega Bio-Tek) following the manufacturer’s protocol. Samples were visualised on an agarose gel again to assure purity (i.e., target length (∼200 bp) bands on agarose gels were visible with minimal to no other bands present).

#### Library preparation and sequencing

Purified PCR products were quantified using a Qubit dsDNA HS Assay kit (Invitrogen) and pooled at equimolar concentration. The pooled library was imaged on a Tape Station 4200 (Agilent) to check the estimated target fragment length. Based on the Tape Station results, the library was cleaned a final time using magnetic beads in a 1:1 ratio with library volume. A unique adapter sequence was ligated to each library using the NEXTFLEX® Rapid DNA-Seq Kit for Illumina (PerkinElmer) following the manufacturer protocol (Appendix 2G). After adapter ligation, the libraries were again imaged on the Tape Station and purified with magnetic beads, this time with a 0.8:1 ratio of beads to sample, as per the NEXTFLEX® Rapid DNA-Seq Kit instructions. Each indexed library and PhiX control were then quantified by qPCR using the NEBNext® Library Quant Kit for Illumina (New England Biolabs). Each library was loaded at a final molarity of 70 pM with a 10% PhiX spike-in. The libraries were sequenced at Liverpool John Moores University on an Illumina iSeq100 using iSeq i1 Reagent v2 (300 cycles).

### Bioinformatic Pipeline

The sequence libraries were demultiplexed by specifying FASTQ generation on the sample run sheets in the Local Run Manager which resulted in forward and reverse FASTQ files for four libraries. All were processed following the same steps, with modifications made based on the primer used. Many functions from the Python (v 2.7.5) package, OBITools (v 1.2.11) ^37^, were implemented in the following pipeline. Commands and functions will be referred to in the format “name of software package”::”function”. First, FastQC (v 0.11.9, https://www.bioinformatics.babraham.ac.uk/projects/fastqc/) software was used to assess per base sequence quality. The forward and reverse sequences were trimmed using OBITools::obicut to lengths between 148 to 150 bp (depending on the fastqc results) such that the average quality score was >36 and that all reads had a maximum length of 150 bp. The trimmed reads were paired using OBITools::illuminapairedend and sorted based on quality score such that paired reads >Q30 were processed. The paired reads were demultiplexed using OBITools::ngsfilter and filtered by length (i.e. elas02 libraries 130 to 210 bp; tele02 libraries 130 to 190 bp). Sequences were dereplicated using OBITools::obiuniq. VSEARCH (v 2.15.2) software ^38^, specifically VSEARCH::uchime_denovo was used to remove chimeras. SWARM (v 3.0.0) was then used to cluster sequences with the fastidious option and d = 1 ^39^. Taxonomic assignment was achieved by using a variety of tools and searching for consensus amongst them. First, global reference databases were generated by using OBITools::ecoPCR for both primer pairs (i.e. tele02 or elas02) against the EMBL nucleotide reference database (r143) specifying all vertebrate sequences (taxid=7742) but excluding human (taxid=9606). Then OBITools::ecotag was used to assign taxonomy referencing the corresponding global databases. Local databases were made using meta-fish-lib to generate custom reference databases ^40^. For the North Atlantic, the latest release of a UK reference database (v258) was downloaded from the github repository ^40^. For the aquarium, a custom species list was created including all species for which a genus was listed in the aquarium inventory followed by executing the meta-fish-lib pipeline ^40^. For each location, a species list was made by supplying the ISO country code for each place: Cape Verde (132), Jordan (400), and South Africa (710) followed by executing the meta-fish-lib pipeline ^40^. For California, a regional 12S reference database called FishCARD was downloaded ^41^. Ten human 12S sequences were added to each local database, anticipating human sequences that are known to be amplified by these primer sets. Taxonomy was then assigned using a few algorithms: 1) OBITools::ecotag, a lowest common ancestor approach against with the corresponding global reference ecoPCR databases and using 2) VSEARCH::sintax, a k-mer approach using a Naïve Bayesian Classifier with the corresponding local reference databases. The BLAST+ suite ^42^ was also used to perform a blastn with 10 maximum targets and a word length = 7 against the local reference databases. Then final taxonomy was assigned as such: 1) Ecotag best identities < 70% were discarded; 2) MOTUs which had a three-way consensus (i.e. ecotag, sintax and blastn returned the same taxonomy) were assigned accordingly; 3) when three-way consensus was not possible, if MOTUs had a two-way consensus between sintax and the blastn results, and both of which were >95% identity then they were assigned accordingly; 4) Remaining MOTUs which did not have consensus between the methods were assigned with the ecotag given assignment since the global vertebrate database was better suited for these cases; 5) The final assignments were filtered by an identity of ≥98%. The final assignments were manually inspected, after the decontamination process explained under the statistical analysis section, to make sure taxonomic assignments made ecological sense. Some changes were made, for instance in the aquarium samples: an inventory list was used to aid in species assignment and some assignments were assumed to be part of the inventory if there were closely related fish that appeared visually similar (Tables S7-S8). Downstream statistical analysis is described in section C of the supplementary methods.

## Results

### Sequencing Overview

A total of 190 samples were sequenced, including 16 syringe-filtered aqueous eDNA samples, 10 metaprobes from the timed soaking experiments and 43 diver metaprobes, most of which were sub-sampled for laboratory protocol testing (Table 1, table S1). The combined sequencing libraries resulted in 6,172,921 reads after bioinformatic quality control, of which 90% (5,579,301 reads) were assigned taxonomy with an identity score ≥ 98% and retained for analysis.

Taxonomic assignment was achieved by using global and local reference databases and searching for consensus between assignment methods (table S2-S6). While most of the samples were collected in the cold-temperate North Atlantic from coastal dive sites in England, Scotland, and Norway (Table 1), seven metaprobes were also contributed from Cabo Verde, the Red Sea (Jordan), the Indian Ocean (South Africa), and the Pacific coast (California) to show-case the global applicability of the approach (table S1). When combining the 39 metaprobes collected by divers from natural ecosystems, 275 unique taxa were identified to either genus or species-level (Fig. 1).

### Testing eDNA capture in an aquarium

To benchmark eDNA capture with metaprobes against the standard method of manual filtration, aquarium staff divers wore metaprobes during two routine dives (Fig. 2). In tandem, we filtered four 1.5 L bottles of seawater for eDNA collected from the same 3.8 million L tropical ocean display exhibit. Additionally, and independent of the divers, metaprobes were soaked to isolate and understand the relationship between time exposed to water and eDNA capture. The metaprobes were suspended in a bleach-washed dive bag, for varying lengths of time up to a maximum of 240 minutes (Fig. 2; fig. S1).

**Fig 2.**
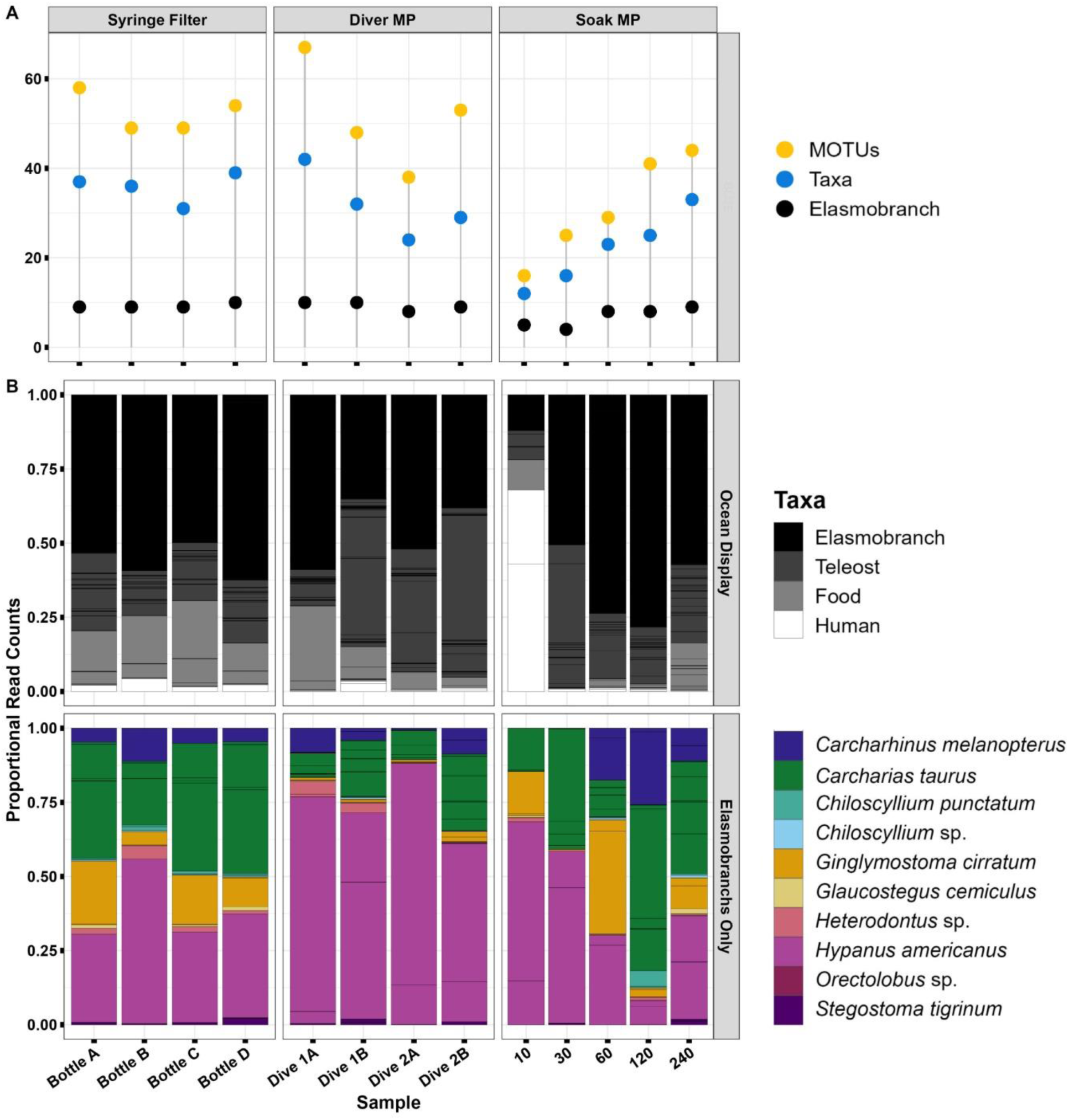
Detections across different sample types in the Ocean display exhibit. **(A)** Lollipop plot showing the number of unique MOTUs, taxa, and Elasmobrachs from each sample. (**B)** Stacked bar charts (top) showing the proportion of MOTUs assigned to Elasmobranchs or Teleosts listed in the Ocean Display inventory. Stacked bar charts (bottom) showing the proportion of MOTUs assigned to Elasmobranch taxa. Sections of each bar separated by black lines indicate unique MOTUs.

A primer designed for Elasmobranch detection was used to analyze the samples from the ocean display, where 12 Elasmobranch species are housed. Only seven of the 12 elasmobranch taxa could be identified to species-level, due to a lack of 12S reference sequences for four of the species and an inability to confidently (≥ 98% identity score) assign *Heterodontus zebra* (Table 2). Despite lacking references sequences for some species, sequences of other species belonging to the same genera were used to assign genus-level taxonomy which resulted in 10 elasmobranch taxa detected overall (i.e., 7 species-level, 3 genus-level) (Fig. 2).

**Table 2.** Inventory of elasmobranch species present in the Ocean display exhibit. Reference sequence availability and detections of the elasmobranchs from eDNA metabarcoding analysis.

| Inventory |  | Reference Sequence Available |  | Detection* |
| --- | --- | --- | --- | --- |
| Scientific name | Common name | species-level | genus-level |  |
| *Carcharhinus melanopterus* | Blacktip Reef Shark | Yes | Yes | Yes |
| *Carcharias taurus* | Sand Tiger Shark | Yes | Yes | Yes |
| *Chiloscyllium arabicum* | Arabian Carpetshark | No | Yes | genus-level |
| *Chiloscyllium punctatum* | Brownbanded Bamboo Shark | Yes | Yes | Yes |
| *Ginglymostoma cirratum* | Nurse Shark | Yes | Yes | Yes |
| *Glaucostegus cemiculus* | Blackchin Guitarshark | Yes | Yes | Yes |
| *Heterodontus portusjacksoni* | Port Jackson Shark | No | Ye s | genus-level |
| *Heterodontus zebra* | Zebra Bullhead Shark | Yes | Yes | genus-level |
| *Hypanus americanus* | Southern Stingray | Yes | Yes | Yes |
| *Orectolobus hutchinsi* | Western Wobbegong | No | Yes | genus-level |
| *Orectolobus maculatus* | Spotted Wobbegong | No | Yes | genus-level |
| *Stegostoma tigrinum* | Zebra Shark | Yes | Yes | Yes |

\*Detection status was the same for syringe-filtered samples and metaprobes worn by SCUBA divers.

Of the 10 elasmobranch taxa detected overall, nine were detected using active filtration in three of the eDNA bottles (i.e., *Orectolobus* sp. missing) but all 10 were detected in Bottle D (Fig. 2A). All 10 elasmobranch taxa were detected in Dive 1 from both samples, but in Dive 2 *Orectolobus* sp. was not detected at all and *Chilosyllium* sp. was not detected in replicate A (Fig. 2). From the soaking experiment, five elasmobranch species were detected in 10 minutes but by 60 minutes all ten of the identifiable elasmobranchs were detected (i.e., when combining all six samples from 10, 30, and 60 minutes). Without combining time points, the samples that soaked for 240 minutes were the only ones that individually contained all 10 elasmobranch taxa (Fig. 2B).

The soaking experiment was replicated in a smaller exhibit, referred to hereafter as the ‘coral cave’, which contained tropical fish species. The teleosts, which were the target taxa of the primer used for metabarcoding of the coral cave samples, also increased in number of detections over time, with the top ten most read abundant taxa detected by 60 minutes soaking time (Fig. S2). Combining data from the coral cave and the ocean display, soaking time was found to be a statistically significant factor determining the number of MOTUs (R^2^ = 0.34, p = 0.004) and unique taxa detected per sample (R^2^ = 0.60, p < 0.001) (Fig. 3, table S9). When adding the samples collected by divers, which were collected from 50-minute and 65-minute dives, the relationship between time and detections weakened, but was still significant (MOTUs: R^2^ = 0.15, p = 0.02 and Taxa: R^2^ = 0.26, p = 0.003) (Fig. S3, table S9).

**Fig 3.**
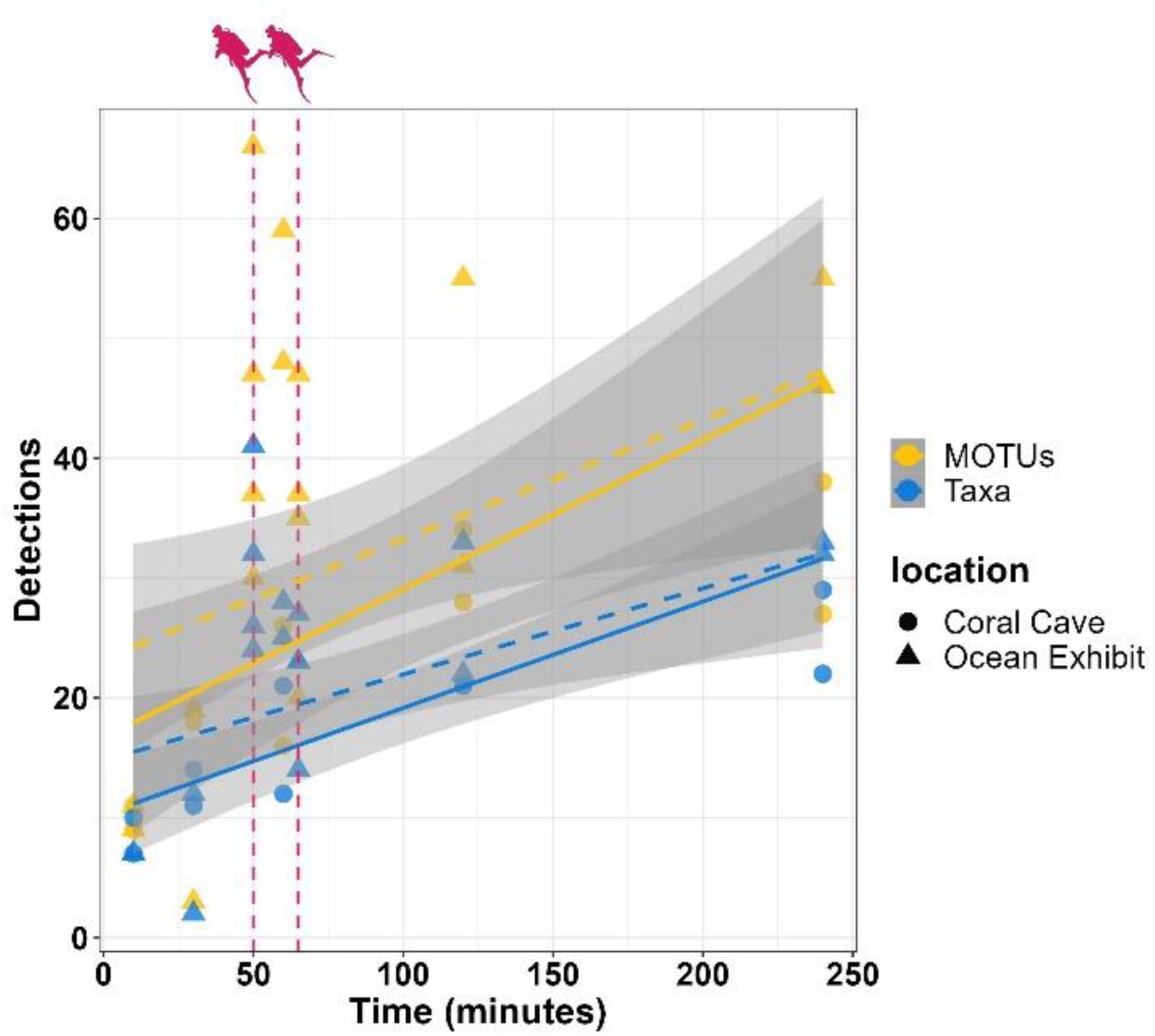
Number of MOTUs and number of taxa detected from metaprobes over the course of the soaking experiments. Samples are combined from the Ocean display and Coral Cave tanks and were collected at the time intervals: 10 min, 30 min, 60 min, 120 min, and 240 min. Linear regression lines are overlayed (solid line). The vertical dashed lines in red represent the diver metaprobes from the Ocean display added in at 50 and 65 min (vertical dashed lines). Dashed regression lines represent the linear regression models recalculated to include dive data.

### Comparing eDNA capture methods

eDNA capture by filtering 1.5 L of water was compared to eDNA collected on metaprobes worn by SCUBA divers at four locations: the Ocean exhibit at the Blue Planet Aquarium, Dukes Dock in Liverpool, and the SMS Bayern and SMS Brummer dive sites in Orkney (Fig. 4). The syringe filtered eDNA from samples collected in nature resulted in a higher amount of unique taxa detections (i.e., between 65-73%) compared to unique taxa detected in metaprobe eDNA (i.e., 4-10%) (Fig. 3B-D). This differed from the samples collected at Blue Planet Aquarium which had a large proportion of taxa (∼65%) detected by both methods, rather than unique taxa from a single method representing a majority (Fig. 4A-D). Regardless of sample type, beta-diversity among dive site locations significantly differed (R^2^ = 0.304, p = 0.001, Fig. 4E, table S10), while beta-diversity did not statistically differ between sample types, when controlling for dive site location (R^2^ = 0.045, p = 0.057, Fig. 4E, table S10).

**Fig 4.**
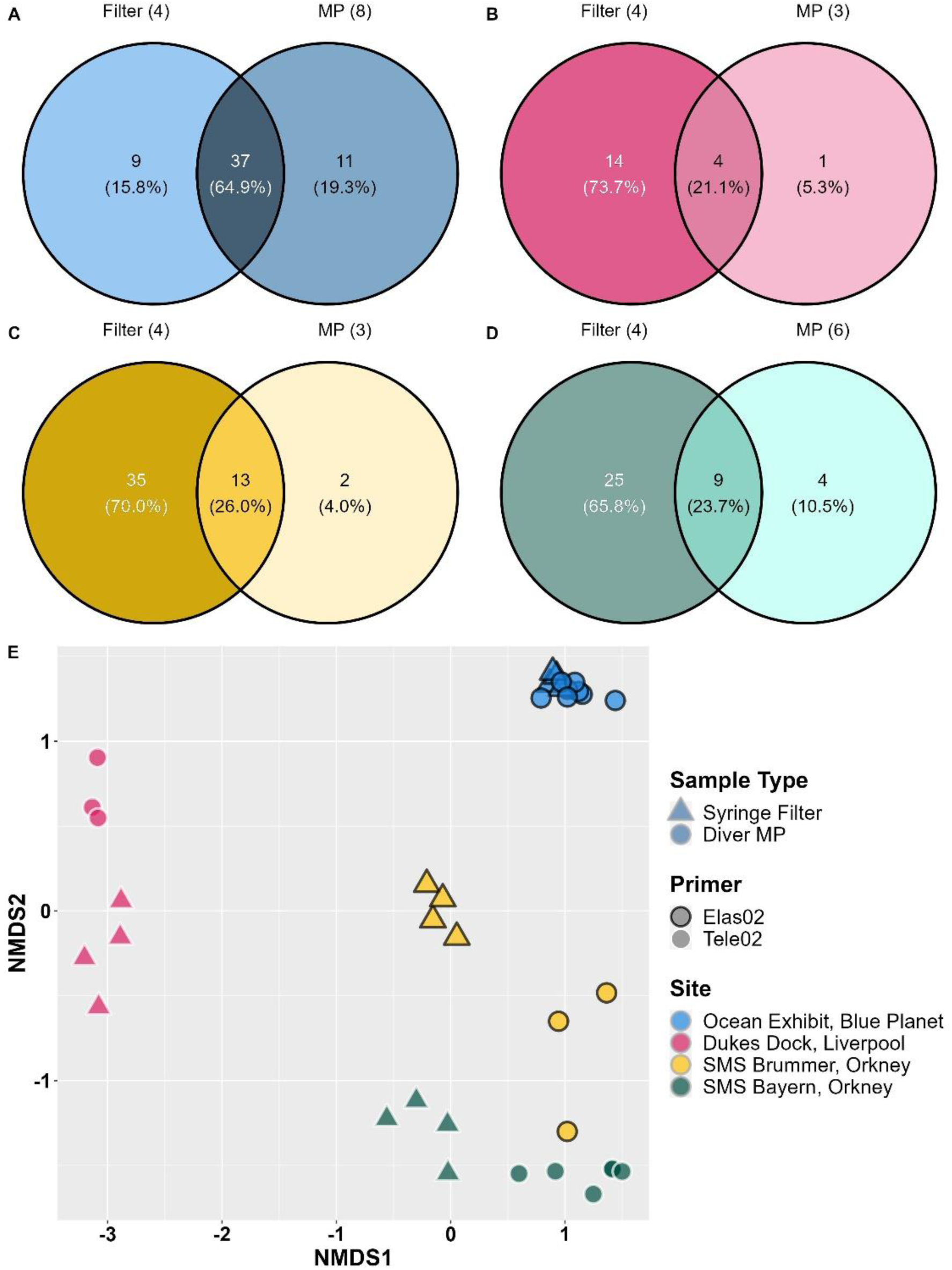
Comparison of detections from syringe-filtered eDNA and metaprobe eDNA. Venn diagrams showing number of genus and species-level taxa detected using syringe filtered water versus divers with metaprobes at **(A)** the Blue Planet Aquarium, **(B)** Dukes Dock in Liverpool, **(C)** SMS Brummer in Orkney and **(D)** SMS Bayern in Orkney. Shading of the sets and unions darken with increasing number of taxa. Sets or unions with the highest number of taxa in each Venn diagram have white text. **(E)** Non-metric dimensional scaling plot of Jaccard distances for genus and species-level taxa from the four sampling events and locations.

### Refining field and laboratory protocols

To evaluate which field and laboratory techniques work best, we first tested whether the amount of input gauze material and lysis buffer affected the outcome of the DNA extractions in terms of species richness, given the uncertainty around the degree of randomness by which DNA adheres to the cotton-gauze. We found no significant difference between input amounts of cotton or lysis buffer (p = 0.704, p = 302, respectively; Table S11; fig. S4). When comparing different DNA extraction methods, we found no significant difference between techniques (i.e., Mu-DNA or Qiagen DNeasy Blood and Tissue kit) (p = 0.671, table S12). Finally, we tested whether the different preservation techniques influenced species richness and found that there was a strong effect of preservation, with ethanol preserved sub-samples being much more likely to have higher species richness than the corresponding silica gel preserved cotton (p = 0.006, Table S13).

### Volunteers scale up eDNA capture locally and globally

In the UK, volunteer efforts provided a snapshot of North Atlantic inshore fish biodiversity (Table 1), resulting in 84 unique genus and species-level taxa detected. The vertebrate communities detected at dive locations were significantly separated according to ICES advisory areas, which are established in part by biogeography but also for fisheries management purposes (p = 0.001, Fig. 5, table S14). Species that are common and widely distributed in the North Atlantic, such as Atlantic cod (*Gadus morhua*), silvery pout (*Gadiculus argenteus*), butterfish (*Pholis gunnellus*), Northern rockling (*Ciliata septentrionalis*), did not cluster near a single set of samples but instead occupied neutral space, indicating their importance across multiple locations.

**Fig 5.**
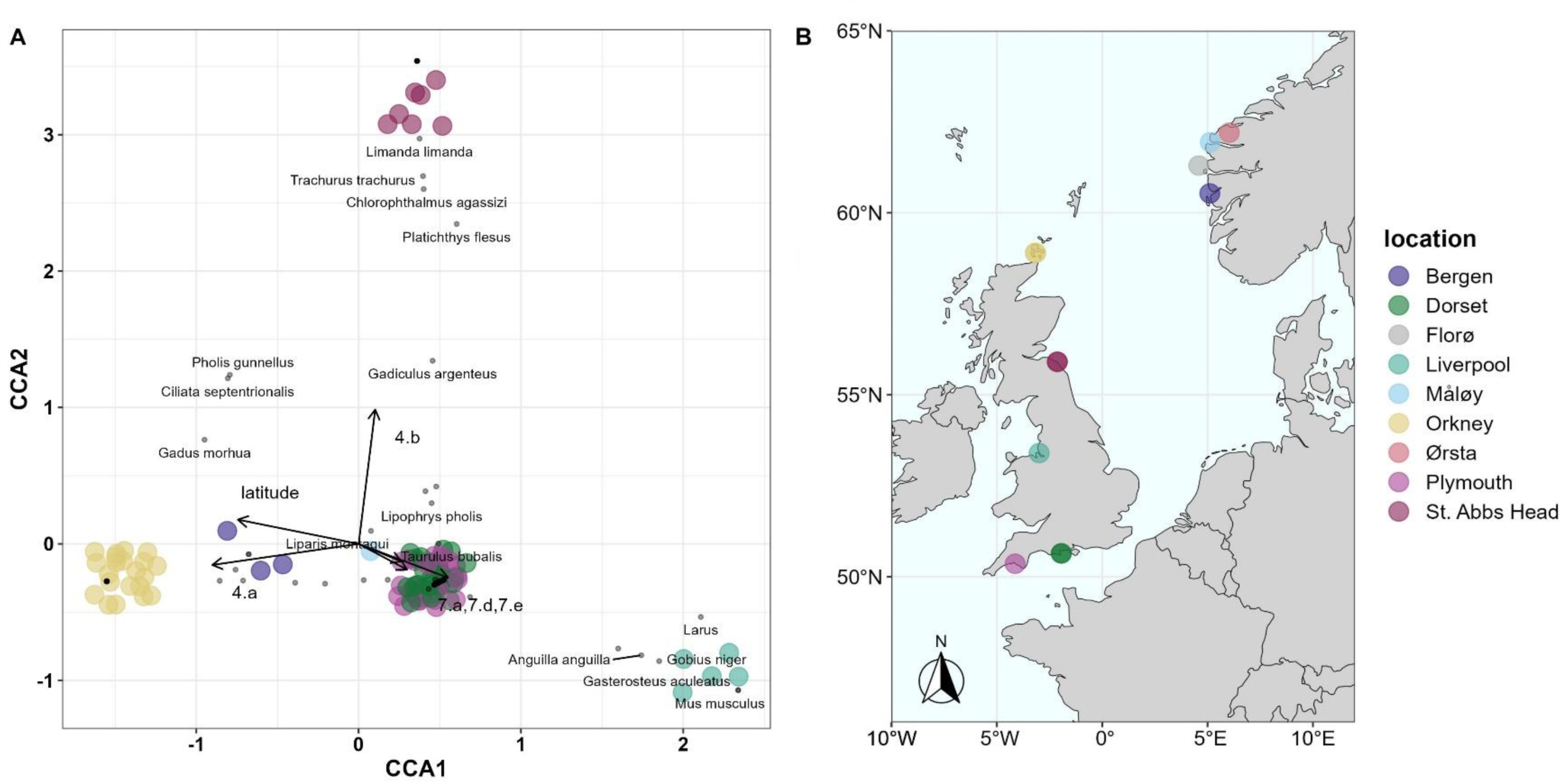
North Atlantic metaprobe samples. **(A)** CCA plot showing how communities detected North Atlantic dive sites separate by ICES advisory areas (4.a, 4.b, 7.a, 7.d, 7.e) and latitude. Location points are randomly jittered by 0.2 for visibility. (**B)** North Atlantic dive site locations. Florø is not included in the CCA plot since the samples (among those preserved in silica beads) failed.

Outside of the UK, we were able to collect samples from six countries in total: Cabo Verde, England and Scotland (United Kingdom), Jordan, Norway, South Africa, and the USA (Table 1). When plotting the samples and species on a CCA ordination, the countries significantly differed along the latitudinal gradient (p = 0.001) and between ocean basins (p = 0.001) (Fig. 6A, table S14). Most species points clustered around corresponding samples from each country, with one clear exception: the cosmopolitan pelagic ocean sunfish (*Mola mola*), which was detected in both the North Atlantic and North Pacific basins (Fig. 6A). Out of 180 species-level detections, 18 (10%) were classified as ‘near threatened’ or more severe, according to the IUCN Red List criteria (Table S15). Two critically endangered fish, the European eel (*Anguilla anguilla*) and the giant guitarfish (*Rhynchobatus djiddensis*) were detected in England and South Africa, respectively (Fig. 6B). An endangered ray, the reticulate whipray (*Himantura uarnak*), was detected in South Africa. In total, there were six Elasmobranchs classed as vulnerable which included for example, the thresher shark (*Alopias vulpinus*) and spurdog or spiny dogfish (*Squalas acanthias*), detected in the USA and Norway, respectively. Of all the countries, samples from South Africa contained the most threatened species detections even though only two dives were performed in the country with only two metaprobes deployed for each, and all of which were preserved in silica beads for ease of transport (Table S15).

**Fig 6.**
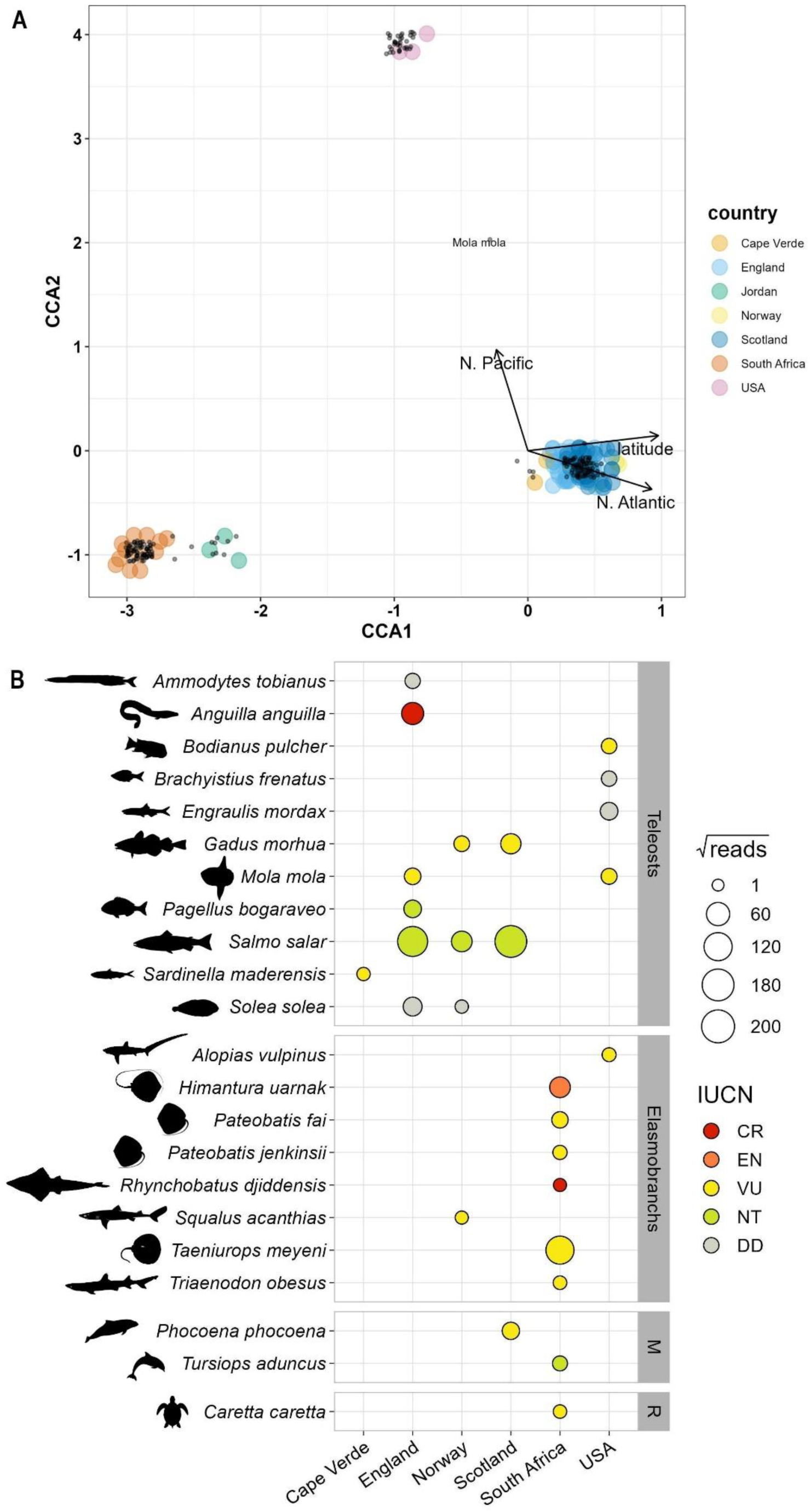
All metaprobe samples and detected threatened species. **(A)** CCA plot showing how communities detected at global dive locations separate by ocean basin and latitude. Location points are randomly jittered by 0.2 for visibility. The inset map shows countries from which samples were collected from in red. **(B)** Bubble plot showing square-root transformed read counts of detections within the following IUCN red list categories: ‘critically endangered’ (CR), ‘endangered’ (EN), ‘vulnerable’ (VU), ‘near-threatened’ (NT) and ‘data deficient’ (DD). Taxa within the ‘least concern’ and ‘not evaluated’ categories are not shown. ‘M’ and ‘R’ are abbreviations for Mammals and Reptiles, respectively.

## Discussion

Monitoring biodiversity is essential for effective stewardship of marine ecosystems and services. Comprehensive monitoring is often not viable due to the diversity of the habitats and life, which can necessitate different and costly approaches ^24^. Environmental DNA (eDNA) analysis is a highly sensitive, non-invasive, and universal approach for different marine taxa, providing extensive species distribution information ^25,26^. Moreover, there are ongoing efforts to implement eDNA collection as a key monitoring strategy in various countries ^14,27,28^. Concentrating eDNA typically involves water filtration, but more recently passive techniques whereby DNA adheres to a surface over time, have been explored. Here we used a quasi-passive eDNA capture technique and a network of snorkeling and SCUBA diving volunteers to collect eDNA from marine environments (Fig. 1).

### Protocol refinement

The sampling protocol was iteratively refined based on differences in sequencing results and on what was most convenient for our sampling purposes. The preservation method (i.e., silica gel beads or 100% ethanol) affected species richness detection such that replicates of samples preserved in ethanol performed significantly better. These metaprobe samples were exposed to the same field and laboratory conditions, but with varying preservation techniques, we therefore divided the cotton gauze and preserved the sub-samples differently to compare the outcome. We did this 18 times (Table S1), and nine (50%) sub-samples that had been preserved in silica beads resulted in no detections of target taxa, whereas all corresponding ethanol preserved sub-samples had detections. Despite preservation by silica gel dehydration yielding unsatisfactory DNA recovery compared to ethanol, the samples collected from Sodwana Bay, South Africa were preserved in silica beads to ease shipping logistics, and 11 out of those 12 samples contained target taxa. Notably, the sampling containers used in South Africa were large (200 ml) and each gauze piece was placed in a separate container (i.e., 3 seperate containers for cotton material from 1 metaprobe), allowing a volumetric ratio of silica beads to sample ∼3:1. It seems that silica beads can work if the cotton gauze is completely dehydrated, but to practically achieve dry conditions with the constraints of materials was difficult. This differed from previous research in which metaprobes were placed in bottom-trawl nets, and no significant difference was found between the efficiency of ethanol and silica gel bead preservation methods ^23^. The commercial extraction protocol (Qiagen Blood and Tissue kit) did not significantly differ from the non-commercial method (Mu-DNA Tissue protocol) ^29^ and the same was true for different amounts of input cotton gauze and lysis buffer. We found that preservation in ethanol and using the Qiagen Blood and Tissue DNA extraction kit (i.e., with 1000 µl of the provided lysis buffer, Buffer ATL, per DNA extraction) with between 0.2-0.4 g of cotton-gauze, produced the most consistent and reliable results.

### Benchmarking against conventional eDNA capture

Diver metaprobes differentiated vertebrate communities from various sampling locations and detected similar communities to corresponding aqueous eDNA samples. In an aquarium setting, diver metaprobes compared to typical filtered aqueous eDNA samples resulted in statistically similar community compositions, where ∼65% of the fish detected were the same. When comparing detections of elasmobranchs, at least one sample from each method contained all 10 of the total unique elasmobranch detections (Fig. 2). The diver metaprobes however, contained more unique detections than did the filtered eDNA (i.e., ∼20% versus ∼15%) (Fig. 4). These observations differed markedly to diver and filtered samples collected in nature, where the overlap in detections by each method was between 20-30%, with filtered eDNA samples containing between 65-75% of unique taxa (Fig. 4A-D). This percentage of overlap was lower compared to a previous study using active filtration and passive eDNA capture on different types of suspended filters where active and passive methods shared between 43%-75% of species detections; however, this was with ∼70 passive capture replicates deployed at each site with soaking times between 4-24 hours ^18^, which is not a feasible scenario with SCUBA diving. The volume of filtered water needed to capture the eDNA necessary for a representative sample of biodiversity can differ depending on the target taxonomic group ^30^ and the habitat being sampled ^31,32^, and similarly we would anticipate that factors influencing exposure to the environment: such as time in the water, distance covered by a diver and site hydrodynamics as well as habitat type, all could profoundly change the optimal sample size. Clearly, our experiments in an aquarium show that detections increased statistically with time, but that this relationship weakened in strength when adding in the samples collected by divers (Fig. 3), which suggests that the swimming motion of a diver can increase the eDNA accumulation rate in the gauze, compared to a stationarily soaking metaprobe. Previous passive eDNA capture field experiments show that species richness did not increase with time ^18^ and that the relationship between submersion time remains unclear ^33^ but it is possible that over longer sampling periods the eDNA has time to desorb (after initial adsorption) from the capture substrate ^19^. Establishing these thresholds for diver metaprobes in the future in targeted habitat types could enable further understanding of how to implement this monitoring strategy. At the four locations where we collected both filtered eDNA samples and diver metaprobes, community structure did not significantly differ between sampling strategies (Fig. 4E). Moreover, when comparing the vertebrate communities detected in temperate North Atlantic habitats (i.e., coastal UK and Norway), the communities at sampling locations could be segregated according to their corresponding ICES advisory area (Fig. 5), demonstrating the capability for this approach to be employed for large-scale monitoring.

### Towards a global effort

One-hundred-forty-six samples were collected in six countries by volunteers, resulting in 275 vertebrate taxa assigned to either a genus or species, 180 of which were identified to the species-level. In some cases, such biodiversity records were obtained by a modest number of metaprobes: samples collected in South Africa came from two dives (i.e., by one dive buddy pair), resulting in cotton material from just four individual metaprobes, while the Cape Verde, California, and Jordan sampling events consisted of only one dive at each, with just one diver of a buddy pair wearing a metaprobe. The cotton from these metaprobes was sub-sampled in triplicate for sequencing, representing just under 15% of the sequenced samples (n = 21), but recovering 132 marine taxa (14 genera, 118 species). Despite the relatively low sampling effort outside of the North Atlantic, vertebrate communities detected at the dive sites significantly differed between countries and grouped by ocean basins and across latitude (Fig. 6A). There were certain vertebrates that were unique to dive sites and made up a large portion of read counts, for example: 17,933 reads of California sea lion (*Zalophus californianus*) in California reflecting the nearby colony, 14,158 reads of round ribbontail ray (*Taeniurops meyeni*) from 9-mile reef in South Africa, and 47,524 reads of three-spined stickleback (*Gasterosteus aculeatus*) from Dukes dock in Liverpool, UK. Round ribbontail rays are one example of the 12 species we detected which are classed as vulnerable by the IUCN (Fig. 6B). With relatively low effort sampling, we detected important fisheries species that are classed as data deficient by the IUCN including lesser sand eels (*Ammodytes tobianus*) and common sole (*Solea solea*). Some detections were of rare elasmobranchs such as the thresher shark (*Alopias vulpinus*) in California and the giant guitarfish (*Rhynchobatus djiddensis*) in South Africa, highlighting how these data collected by volunteers could meaningfully contribute to global biodiversity data repositories.

This project, conveyed as ‘DNA divers’ to the general public, presents an example of how marine recreationists can be engaged with eDNA studies and in doing so learn about emerging technologies and the importance of monitoring biodiversity more broadly. There have been successful citizen science programs working with volunteers either to collect eDNA or to go SCUBA diving for visual surveys. Metaprobes can be 3D printed, cotton medical gauze is widely available, and these materials can be easily assembled and attached to a diver’s BCD (buoyancy control device) or weight belt with cable ties. This has the advantage of there being little to no training required for SCUBA divers or snorkelers to participate in collecting marine eDNA, relative to visual census, which requires taxonomic training. Moreover, in the UK, visual census undertaken by recreational divers cannot be executed with industry standard survey methods, such as transects, since this violates The Diving for Work Regulations 1997. This means that any eDNA metabarcoding data collected by volunteers is concomitant with visual census information and could provide two-layers of species presence data associated with the dive site more generally and without violating diving for work laws. At Dukes Dock in Liverpool, volunteer Seasearch divers completed a visual census survey (Fig. S7) while collecting eDNA that was ultimately screened for teleost species; both methods contained observations of three-spined stickleback (*Gasterosteus aculeatus*) and black goby (*Gobius niger*). The divers recorded all types of taxa they could see, including invertebrates like mussels (*Mytilus edulis*), European green crab (*Carcinus maenus*) and yellow sun sponge (*Halichondria bowerbanki*), while the metaprobe eDNA was analysed targeting a vertebrate gene therefore capturing urban landscape-level species too, like gulls (*Larus* sp.) and house mice (*Mus musculus*). The second most sequence read abundant fish was the critically endangered European eel (*Anguilla anguilla*), unseen by the divers, but divers saw two-spotted goby (*Gobiusculus flavescens*), undetected in the eDNA. Visual observations will always remain a valuable source of marine biodiversity data, but passive eDNA collection could be a way to crowdsource molecular samples, supplement monitoring efforts, and include a wider demographic of SCUBA divers and marine leisure users (i.e., kayakers, paddle boarders) who either do not have the time, interest, or resources to engage in taxonomic training.

There is broad interest in passive methods for non-invasive biodiversity surveillance using eDNA metabarcoding ^19–22,33,34^. The semantics of what constitutes ‘passive surveillance’ can differ. Some methods have been labelled as passive due to the automated process of water filtration using marine robotics ^5,35^, while other passive techniques have explored DNA binding to submerged materials ^19,20,22^. In our case, we rely on the latter as well as the exertion and exploration activities of volunteer SCUBA divers and snorkelers. This study benefitted from the involvement of ∼30 divers – some of whom are co-authors of this study – who went on dives in four continents across a range of habitats (i.e., seagrass beds, kelp forests, shipwrecks, fjords, coral reefs), resulting in the detection of 275 different taxa overall. Although the diver metaprobes likely capture fewer species than what it is usually possible using traditional water filtering from a given area, they still proved effective at significantly distinguishing assemblages at regional and global scales and detecting species of conservation importance with relatively low effort. While we are not suggesting that our method excludes the need for engineering autonomous eDNA sampling instruments, the vast potential of the ‘DNA divers’ approach is undeniable: many coastal marine areas remain poorly monitored, while volunteers seem to be very keen to embrace this method. Through such a cost-effective approach, it is reasonable to expect that eDNA samples from virtually every region of the world can be rapidly amassed, thereby upscaling and boosting marine monitoring, and at the same time raising public awareness and improving science literacy.

## Acknowledgements

We would like to thank Ivvett Modinou for her unwavering support, belief, and guidance with public engagement materials. We are grateful to Ainsley Hatt for connecting us with the British diving community. We are indebted to all volunteers who participated in the DNA divers project, both on the authorship team or listed here: Lisa Shafe, Michael Moore, Cath Lee, Racheal Priest, Carolyn Waddell, Donovan Lewis, Rory McWilliams, Mike Cliff, Sophia Panuthos Buell, Angus Jackson, and others who chose to remain anonymous. We are grateful to Thomas Bryne and the LJMU advanced manufacturing lab for assistance with the CAD 3D-printing design and printers. We acknowledge the generosity of Seasearch coordinators, Halton Charters Ltd., Natural England, Newcastle University sub-aqua club, Sharklife conservation group, and the Blue Planet Aquarium.

## Funding

LJMU QR Policy Support fund to EFN and SM.

LJMU QR Participatory Research fund to EFN and SM.

Natural England commissioned report 506 to EFN and SM.

UK Natural Environment Research Council Grant NE/T007028/1 (SpongeDNA) to SM and Ana Riesgo.

