## supplementary for "DNA Divers: volunteer-based eDNA capture for local and global marine biodiversity monitoring"

### Supplementary Methods

#### A. Sample Collection (supplement)

***Metaprobe Material Preparation and Field Sampling Methods***

These methods have been written for a non-technical audience and provide instructions for how volunteers can prepare metaprobes. The cotton gauze (~ 10 cm x 3.7 m) should be certified sterile and can be found on various medical supply websites. The same methods were followed if metaprobes were prepared in a laboratory, except that after preparation they were treated with 30 min of UV.

***Preparing Metaprobes***

**1.** Rinse dirty metaprobes with tap water, removing any residue or salt. (If not newly printed)

**2a**. Prepare a bleach solution to clean the dirty metaprobes and dirty scissors. Wearing medical gloves, fill one wash basin with a solution made up of 2 parts bleach cleaning chemical and 3 parts bottled drinking water. The depth of the solution should be enough to submerge at least one half of a metaprobe. This bleach solution is ideally 10% bleach, however most household cleaning equipment will not be this powerful. Aim to make the bleach solution strong enough to smell if you are standing directly next to it. Proceed to step 3a.

**2b.** If it is not possible to do step 2a, follow this alternate step. Locate paper towels and a cleaning chemical containing bleach. Proceed to step 3b.

**3a.** Soak the dirty metaprobes and scissors in the bleach solution. The metaprobes should soak for at least half an hour (30 min.). They can be left overnight. The scissors should soak for ~10 min. since the steel will rust if left too long. Proceed to step 4.

**3b.** Spray the dirty metaprobes and scissors with the cleaning chemical containing bleach. Wearing gloves, generously apply bleach cleaning chemical to the metaprobes and scissors. Wipe both dry with paper towels. Repeat three times. Proceed to step 4.

**4.** Prepare a soapy solution to rinse the metaprobes and scissors. Wearing medical gloves, fill the second wash basin with a solution made up of 1 part washing up detergent and 20 parts bottled drinking water. Aim to make a soapy solution that is milder (i.e. not too many bubbles) than what you would use to do the washing up. Transfer the bleached metaprobes and scissors to the soapy wash bath for 5 min. Top tip: Since bleach degrades DNA (cleans the metaprobes) it is critical to remove excess bleach before preparing clean metaprobes. This is an important step!

**5.** Allow the scissors and metaprobe to air dry on paper towels or a clean surface.

**6.** Prepare the cotton gauze for the metaprobes. Clean a table or surface with a cleaning product that contains bleach. Wearing gloves, open the gauze and cut it into quarters. Place three of the quarters into a metaprobe half. The quarters should be about the size of a cotton ball (i.e., large enough so that the gauze is too large to fit through the perforations in the metaprobe). Note that the purpose of pre-cutting the gauze is so that they will fit into sample collection tubes. Depending on the size of your sample collection tubes, this may not be an issue, in which case the gauze roll can remain intact and placed directly into the metaprobe.

Important Note: After trial and error, volunteers found that two whole rolls (10 cm x 3.7 m) of cotton gauze fit into the metaprobe and did not need to be altered. Larger sample collection containers were used to accommodate this. See supplementary table, column 'Gau', indicating whether whole or partial rolls of cotton were used.

**7.** Once the pieces of gauze are in the metaprobes use two to three cable ties to join the opposite halves together.

**8.** Keep the clean metaprobes in a clean storage place until your next dive. This could be a fresh resealable bag or a wash basin that has been wiped clean with bleach cleaning chemicals. Top tip: Make sure this storage area is protected from any seawater spraying into the boat.

***Preserving Metaprobe Samples***

**1.** Health and safety is the #1 priority. Please make sure you have safely exited the water and are in a comfortable position on your boat or on land.

**2.** Preserve the samples within half an hour (30 min.) of completing your dive.

**3.** Cut the metaprobe free from your dive equipment. Cut the cable tie attaching your metaprobe to your BCD or other equipment. Store the metaprobe in a clean area. If necessary, a resealable bag can be used to keep the metaprobe clean while you take off your dive gear. Top tip: Ask someone wearing medical gloves to assist you with this step. Medical gloves can be difficult to put on wet hands.

**4.** Wear medical gloves, if not already doing so.

**5.** Cut the metaprobe open on a clean surface. Do not use a surface that is normally used to gut or clean fish. Clean a table or surface with a cleaning product that contains bleach. Cut the cable ties on the exterior of the metaprobe which hold it closed. Top tip: Place the halves of the metaprobe open-side facing up onto the surface. This prevents the gauze from touching the table and helps prevent sample contamination.
**6.** Place each medical dressing roll into a plastic screw-cap tube. The screw-cap tubes will contain either silica beads or ethanol. If silica beads, shake the tube so that the beads surround the gauze. The beads will turn green as they adsorb water. If using ethanol, make sure the sample is submerged. Note the number on the tube and cap of the plastic screw-cap tubes. This number will be recorded on the Collection Log Form.
**7.** Keep the samples in a cool, safe place. Keep the sample tubes in a cold, dark, safe place. Suggested places (in order from best to acceptable): freezer, fridge, cool box, in a plastic bag shieled from sunlight. Once on land, it is highly preferable to keep the samples in a freezer or fridge.
**8.** Fill out the Collection Log Form. A collection log form should be provided with your DNA Divers sampling materials but it can also be found online.

### B. DNA extraction (supplement)

***Qiagen DNeasy Blood and Tissue Kit Method with Modifications***
The following methods have been modified from the ‘Purification of Total DNA from Animal Tissues (Spin-Column Protocol)’ in the Qiagen DNeasy Blood and Tissue Handbook published in July 2020. Unless otherwise stated, the manufacturer protocol should be followed.
**1.** Warm Buffer ATL and Buffer AL to 56°C to fully dissolve any precipitates that may have formed during storage.
**2.** Prepare input material for lysis:
a. For eDNA filters: Using plyers, break open the plastic filter chamber. Over a petri dish, use dissecting scissors and tweezers cut the filter into small pieces. Place half of the filter pieces in a 1.5 ml Eppendorf tube for lysis, and the other half in a 1.5 ml Eppendorf tube for archive at -20°C.

b. For metaprobe gauze: Cut away small pieces of gauze with dissecting scissors, taking sections from various parts. Using tweezers and blotting paper, blot the gauze, changing blotting paper twice, or until most of the ethanol is gone. Weigh the gauze using a weigh boat and adjust the input to between 0.2 g and 0.4 g. Place gauze into a 1.5 ml Eppendorf tube.

**3.** Add 720 mL Buffer ATL and 80 mL Proteinase K, which can be premixed for the number of sample extracts accordingly.

**4.** Mix thoroughly by pulse-vortexing for 5–10 s, and incubate at 56°C in a thermomixer overnight (~16 hours).

**5.** For eDNA filters, centrifuge at 10,000 xg for 1 min at room temperature. This step can be skipped for metaprobe gauzes as they do not form a pellet.

**6.** For eDNA filters, without disturbing the pellet, transfer the supernatant to a fresh 1.5 mL Eppendorf tube. Using the pipette tip, press the gauze to the side of the tube and transfer the supernatant to a fresh 1.5 mL Eppendorf tube.

**7.** Measure the volume of the supernatant and add the same volume of Buffer AL to the sample. Mix thoroughly by pulse-vortexing. Then add the same volume of 100% ethanol. Mix again by pule-vortexing. (e.g., 600 µl supernatant requires the addition of 600 µl buffer AL and then 600 µl ethanol)

**8.** Pipet the mixture into the DNeasy Mini spin column placed in a 2 ml collection tube. Centrifuge at ≥6000 x g (8000 rpm) for 1 min.

**9.** Empty the collection tube and repeat step 8 until all of the mixture has been passed through the spin column. Then replace the collection tube with a fresh tube.

**10.** Add 500 µl Buffer AW1, and centrifuge for 1 min at ≥6000 x g (8000 rpm). Discard flow-through and collection tube.

**11.** Add 500 µl Buffer AW2, and centrifuge for 3 min at 20,000 x g (14,000 rpm) to dry the DNeasy membrane. Discard flow-through and collection tube.

**12.** Place the DNeasy Mini spin column in a clean 1.5 mL or 2 mL Eppendorf tube, and pipet 100 µl Buffer AE directly onto the DNeasy membrane.

**13.** Incubate at room temperature for 1 min, and then centrifuge for 1 min at ≥ 6000 x g (8000 rpm) to elute.

**14.** Pipette the same 100 µl Buffer AE back onto the membrane and repeat step 13 to increase the final DNA concentration in the eluate.

***Mu-DNA Method Reagents***

The following methods are taken from the 'guidelines' section of 'Mu-DNA: a modular universal DNA extraction method adaptable for a wide range of sample types V.2' document on protocols.io (dx.doi.org/10.17504/protocols.io.qn9dvh6).

*Stock solutions*

Stock solutions are given as compositions for 100 mL with the exception of PK.

• 1 M Tris HCl (pH 8):

Dissolve 15.7 g of Tris HCl in 75 mL ddH2O. Adjust to pH 8 with 5 M NaOH. Bring to 100 mL with ddH2O.

• 0.5 M EDTA (pH 8):

Dissolve 18.6 g of disodium EDTA dihydrate in 75 mL ddH2O. Adjust to pH 8 with 5 M NaOH. Bring to 100 mL with ddH2O.

• 20% SDS:

Dissolve 20 g sodium dodecyl sulphate in 75 mL ddH2O, bring to 100 mL with ddH2O.

• Proteinase K (PK)*

*The PK solution described in the protocol was not used. Instead, Proteinase K Solution (20 mg/mL), RNA grade (Invitrogen) was used without modification.

• 5 M Ammonium acetate:

Dissolve 38.6 g ammonium acetate in 75 mL ddH2O, bring to 100 mL with ddH2O.

• 180 mM Aluminium etc.:

Dissolve 8.2 g aluminium ammonium sulphate dodecahydrate in 75 mL ddH2O, bring to 100 mL with ddH2O.

• 3% Calcium chloride:

Dissolve 3 g calcium chloride dihydrate in 75 mL ddH2O, bring to 100 mL with ddH2O.

• 5.5 M Guanidine HCl:

Dissolve 52.6 g guanidine hydrochloride in 75 mL ddH2O, bring to 100 mL with ddH2O.

*Working solutions*

All working solutions are composites of stock solutions. All working solution compositions are given for a 100 mL final volume. The same ratios could be maintained to adjust for different desired final volumes. Note that some working solutions consist of a single stock solution.

• Lysis Solution:

To 75 mL ddH2O add 6.7 mL 1 M Tris HCl (pH 8), 5.3 mL 0.5 M EDTA (pH 8), 1.7 g guanidine thiocyanate, 8.7 g trisodium phosphate dodecahydrate and 0.2 g sodium chloride. Stir mixture until all solids dissolve. Adjust to pH 9.0 with 5 M HCl. Bring to final 100 mL volume with ddH2O.

• Tissue Lysis Additive:

20% SDS

• Flocculant Solution:

To 50 mL 5 M Ammonium acetate add 25 mL 180 mM Aluminium etc. Vortex briefly before adding 25 mL 3% Calcium chloride. Vortex briefly to mix.

• Tissue Binding Solution:

To 50 mL 5.5 M Guanidine HCl add 50 mL 100% ethanol. Vortex briefly to mix.

• Wash Solution:

To 20 mL ddH2O add 80 mL 100% ethanol.

• Elution Buffer:

To 75 mL ddH2O add 1 mL 1 M Tris HCl (pH 8) and 0.2 mL 0.5 M EDTA (pH 8). Bring to 100 mL with ddH2O.

***Mu-DNA Method***

The following methods are a combination of the 'Tissue' and 'Water' protocols adapted from the 'Mu-DNA: a modular universal DNA extraction method adaptable for a wide range of sample types V.2' document on protocols.io (dx.doi.org/10.17504/protocols.io.qn9dvh6).

*Lysis*

**1.** Incubate the Tissue Lysis Additive and Tissue Binding Solution at 55°C to prevent any precipitates that may have formed and until use.

**2.** Prepare input material for lysis:

a. For eDNA filters: Using plyers, break open the plastic filter chamber. Over a petri dish, use dissecting scissors and tweezers cut the filter into small pieces. Place half of the filter pieces in a 1.5 ml Eppendorf tube for lysis, and the other half in a 1.5 ml Eppendorf tube for archive at -20°C.

b. For metaprobe gauze: Cut away small pieces of gauze with dissecting scissors, taking sections from various parts. Using tweezers and blotting paper, blot the gauze, changing blotting paper twice, or until most of the ethanol is gone. Weigh the gauze using a weigh boat and adjust the input amount for the following weight categories: Heavy: 0.9 – 1.1 g; Medium: 0.6 – 0.8 g; and Standard: 0.2 – 0.4 g. Place Heavy amounts into a 50 ml Falcon tube; Medium amounts into a 5 ml Eppendorf tube; and Standard amounts into a 1.5 ml Eppendorf tube.

**3.** Add lysis solution master mix to input material. A lysis solution master mix was made by mixing in 13:1:1 ratio, Lysis Solution (13): Tissue lysis additive (1): Protienase-K (1). 1000 µL of the lysis solution master mix was added to eDNA filters and standard gauze weights. 3000 µL of the lysis solution master mix was added to medium gauze weights. 5000 µL of the lysis solution master mix was added to heavy gauze weights.

**4.** Vortex the tubes and incubate at 55°C for ~16 hours overnight.

**5.** For eDNA filters, centrifuge at 10,000 xg for 1 min at room temperature. This step can be skipped for metaprobe gauzes as they do not form a pellet.

**6.** For eDNA filters, without disturbing the pellet, transfer the supernatant to a fresh 1.5 mL Eppendorf tube. Using the pipette tip, press the gauze to the side of the tube and transfer the supernatant to a fresh tube.

*Inhibitor Removal*

**7.** Add 0.3 X volume of Flocculant Solution (e.g., if 700 µL of lysis supernatant is transferred then 210 µL of Flocculant Solution should be added). Vortex briefly and incubate at 4°C for 10 minutes.

**8.** Centrifuge at 10,000 xg for 1 min at room temperature.

**9.** Without disturbing the pellet, transfer the supernatant to a fresh tube.

*Silica Binding*

**10.** Add 2 X volume Tissue Binding Solution (e.g., if 700 µL of solution from the inhibitor removal step is transferred then 1400 µL of Tissue Binding Solution should be added for a total volume of 2100 µL). Vortex briefly to mix.

**11.** Transfer 700 µL of the mixture to a spin column.

**12.** Centrifuge at ≥ 10,000 xg for 1 min at room temperature, discard the flow-through.

**13.** To standardize the extractions and for practical purposes repeat steps 11 and 12 as follows: a maximum of three times for eDNA filters and standard gauze amounts, exactly four times for medium gauze amounts and exactly five times for heavy gauze amounts.

*Wash*

**14.** Add 500 µL of Wash Solution to the spin column.
**15.** Centrifuge at 10,000 xg for 1 min at room temperature. Discard the flow-through.
**16.** Repeat steps 1 and 2 a second time.
**17.** Centrifuge at 10,000 xg for 2 min at room temperature, replace collection tube with a fresh 1.5 mL Eppendorf tube.

*Elution*

**18.** Add 100 µL of Elution Buffer directly to the spin column membrane and incubate for 1 min at room temperature.
**19.** Centrifuge at 10,000 xg for 1 min at room temperature.
**20.** Take the product in the tube and pipette it onto the spin column membrane. Repeat step 19.
**21.** The DNA is now in the Eppendorf tube.

#### C. Statistical Analysis

Further data processing and statistical analysis were done using R (v.4.1.3). The results of each sequencing library were further decontaminated using the R package decontam (v.1.14.0) [43], specifically by screening the negative controls using the function decontam::isContaminant with the "prevalence" method set to a threshold of 0.5. To control for tag jumping, 0.001% of the total reads for each library was calculated, and any detection lower than 0.001% of the total library reads was removed (i.e. ranging from 2-6 reads). After these thresholds were employed, a small number of species needed to be removed from samples since they did not make ecological sense: *Galeus melastomus* detected in Dorset seagrass beds, *Chimaera monstrosa* a deep-sea species detected in multiple samples (both species could have been laboratory contamination from another project processed at a similar time [44]), *Salmo salar* was detected in California likely because it's a commonly consumed species, *Carcharhinus melanopterus* was an aquarium inventory species in an Orkney sample, and *Carcharias taurus* and *Gymnothorax kidako* were aquarium inventory species that were both removed from St.Abbs samples. Due to the unnatural biomass in an aquarium, the aquarium samples had higher amounts of DNA which could have caused this contamination despite the use of DNA extraction blanks. Finally, from all samples, common domestic species: chicken (*Gallus gallus*), sheep (*Ovis aries*), cow (*Bos taurus*), pig (*Sus scrofa*), dog (*Canis lupus*) and cat (*Felis catus*) were removed.

***Aquarium samples***

Using the inventories provided by the aquarium, taxa detections were classed as either i) inventory species, ii) species used as food, and iii) contamination (Tables S7, S8). Most detections in the aquarium samples were accounted for. For instance, Dalmatian pelican (*Pelecanus crispus*) was removed from the data, but it was likely a true detection as *Pelecanus crispus* is on display in a different part of the aquarium. However, Atlantic salmon (*Salmo salar*) was detected but not listed by the aquarium as a food item nor was it present in an exhibit; therefore, it was classified as contamination and removed from aquarium samples. Using the R package stats (v.4.1.3), linear models (stats::lm) were calculated to assess the relationship between MOTUs and detected taxa over time for both soaking experiments as well as the for the dives in the ocean exhibit display.

***Comparisons of Diver metaprobes to filtered eDNA samples***

To compare the vertebrate communities detected in eDNA collected by filtration versus that of diver metaprobes, various functions from the R packages ggvenn (v.0.1.10) and vegan (v.2.6.4) were used [45]. Venn diagrams were drawn to compare species overlap and uniqueness between sample types. Multivariate homogeneity of group dispersions (vegan::betadisper) was tested for both dive site location and sample type followed by testing for significance (vegan::permutest).

PERmutational Multivariate ANalysis Of VAriance (PERMANOVA) was performed using vegan::adonis2 with the stata parameter set to the dive site location, so that each location was independently considered when comparing the differences in sample type. A second PERMANOVA was carried out to also test for differences in communities by location, independently considering sample type by setting the strata parameter to type.

***Comparisons of preservation, processing and DNA extraction methods***

We tested whether the input weight of cotton gauze effected the alpha-diversity by analysing the species richness and Shannon index in a selection of samples (n=18), each extraction belonging to a different weight category: high (0.9 – 1.1 g, requiring 5000 µl of lysis buffer and five centrifugations), medium (0.6 – 0.8 g, requiring 3000 µl of lysis buffer and four centrifugations), and standard (0.2 – 0.4 g, requiring 1000 µl of lysis buffer and three centrifugations). We used a Generalized Linear mixed model (GLMM) implemented with the R packages lme4 (v.1.1.32) and lmerTest (v.3.1.3), specifically the lme4::lemr function. GLMM was also used to test for the effect of different DNA extraction methods (i.e., Mu-DNA or Qiagen DNeasy Blood and Tissue kit) and to test for the effect of preservation strategy on species richness.

***Differentiating dive sites and locations using CCA model building***

A model-building process was employed to build a minimum adequate model from metadata variables for a canonical correspondence analysis (CCA). The species matrix of square-root transformed read counts was used as a predictor of the CCA. First, a scoping model was made using all of the metadata variables, by setting the intercept parameter in vegan::cca to '.'. Then a blank model was made by setting the intercept parameter to '1'. The function stats::add1 was used to add metadata variables to the model in a stepwise manner, where the object of the function was the blank model, the scope was set to the scoping model, and the test was set to 'permutation'. This stepwise process was repeated until no more variables were significant and an ANOVA (baseR::anova) test was run on the final CCA model to test the strength of the significance for the final model terms. This process was repeated for a subset of the data, specifically samples from the North Atlantic, for which more metadata variables were available.

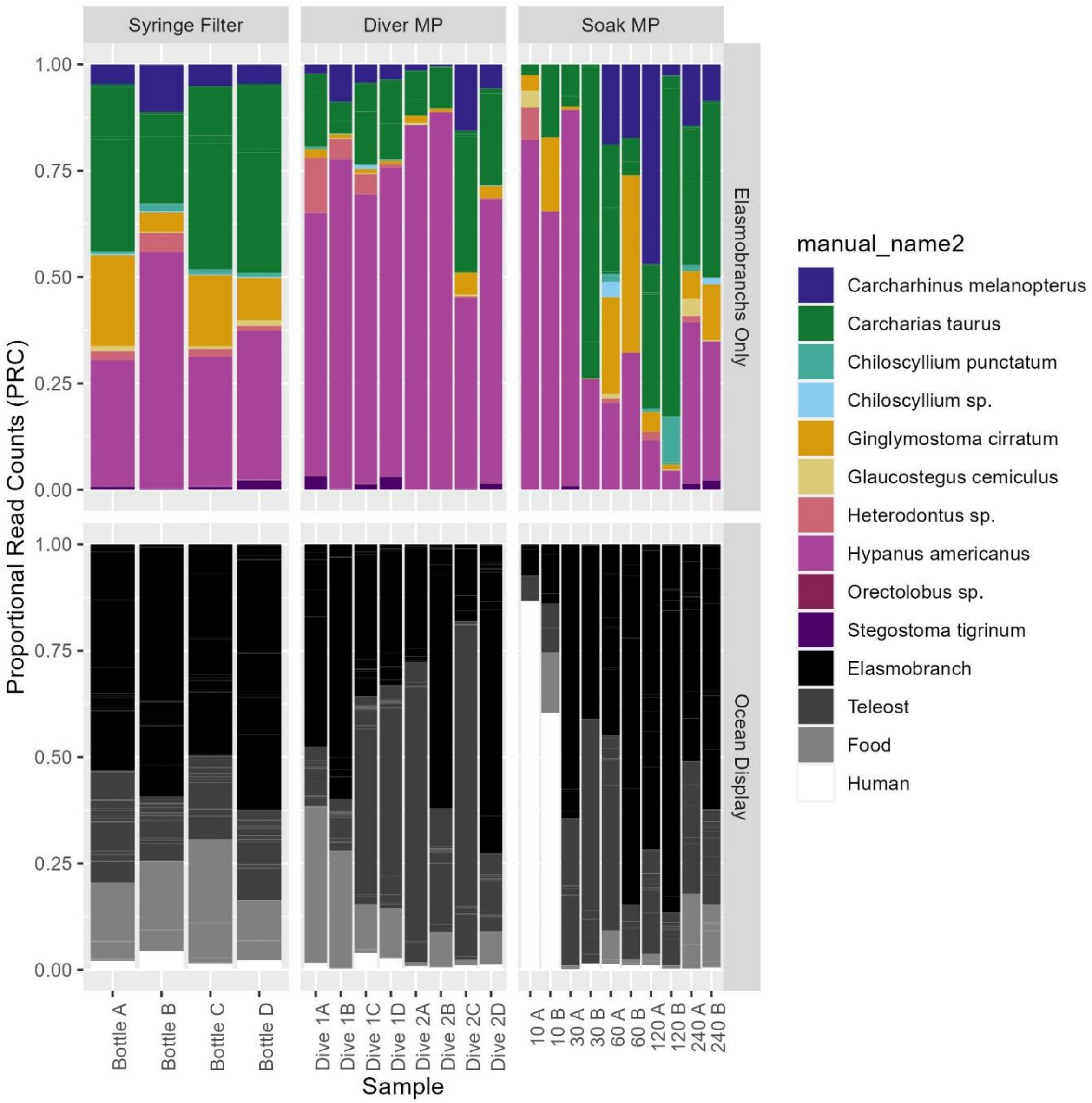

**Fig. S1.**

Stacked bar charts (top) showing the proportion of MOTUs assigned to Elasmobranch taxa with sequencing replicates separated. Stacked bar charts (bottom) showing the proportion of MOTUs assigned to Elasmobranchs or Teleosts listed in the Ocean Exhibit inventory with sequencing replicates separated.

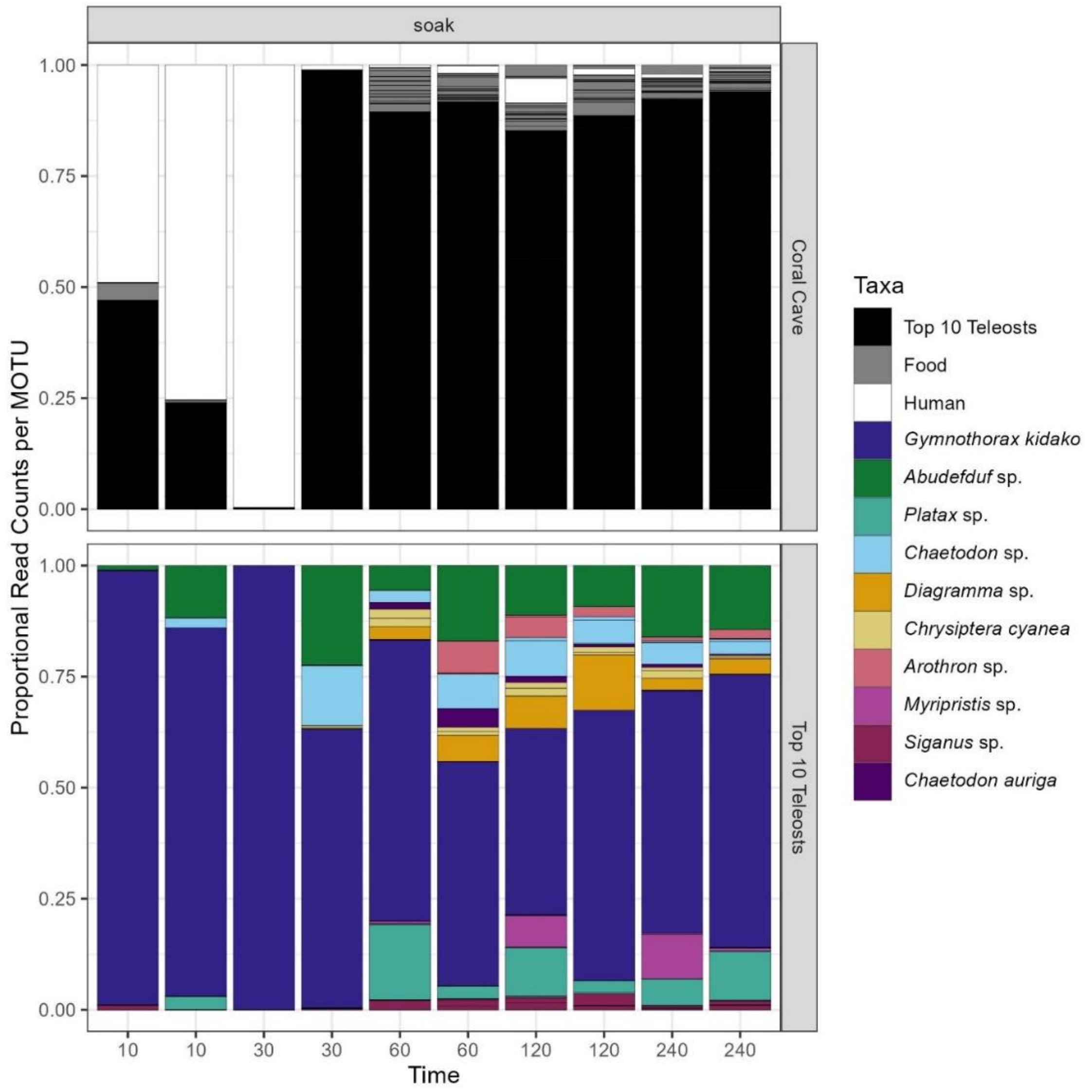

**Fig. S2.**

Stacked bar charts (top) showing the proportion of MOTUs assigned to the top 10 most read abundant Teleosts from Coral cave exhbit with sequencing replicates separated. Stacked bar charts (top) showing the proportion of MOTUs assigned to the top 10 most read abundant Teleost taxa with sequencing replicates separated.

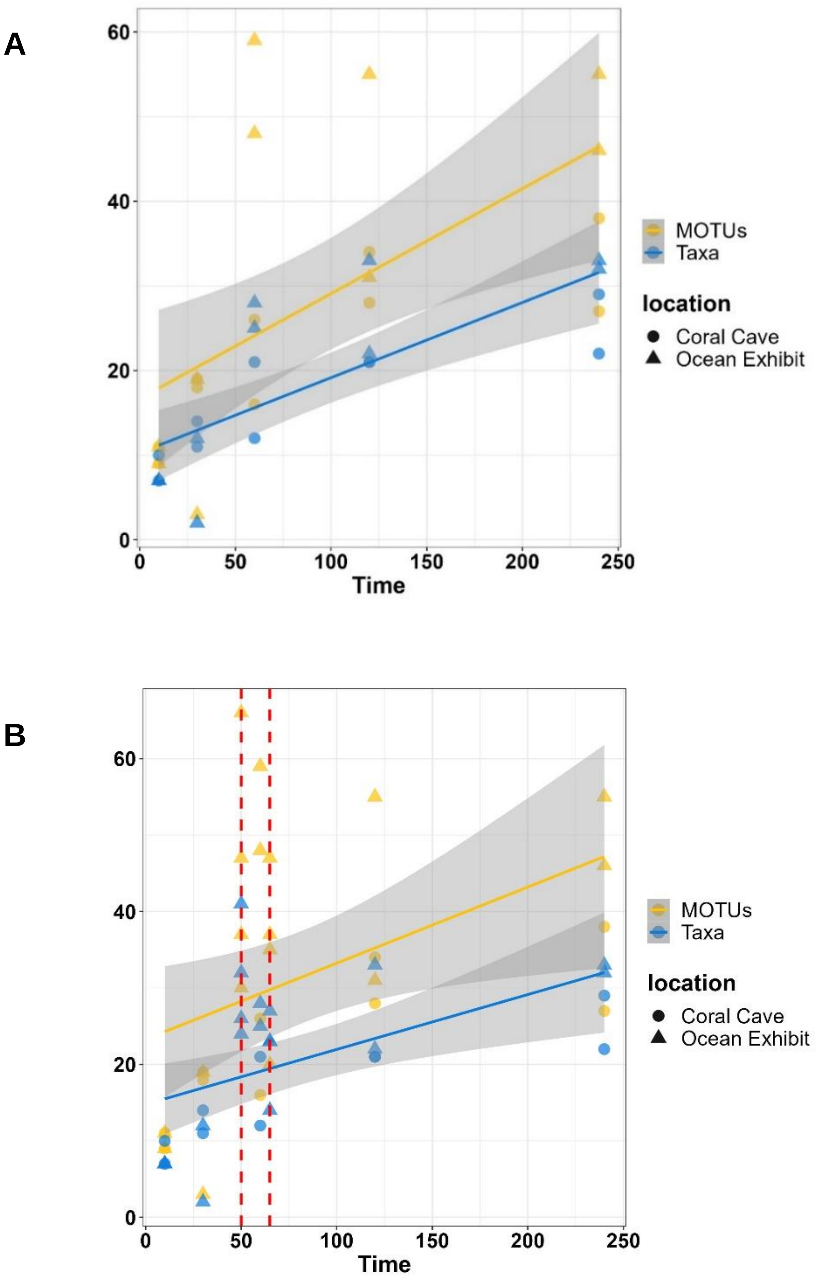

**Fig. S3.**

**(A)** Number of MOTUs and number of taxa detected from metaprobes over the course of the soaking experiments (from the Ocean Exhibit and Coral Cave) at the following time intervals: 10 min, 30 min, 60 min, 120 min, and 240 min. Linear regression lines overlayed. (**B)** Same plot as S3A, with diver metaprobes from the Ocean Exhibit added in at 50 min and 65 min. Linear regression models recalculated to include dives and lines overlayed.

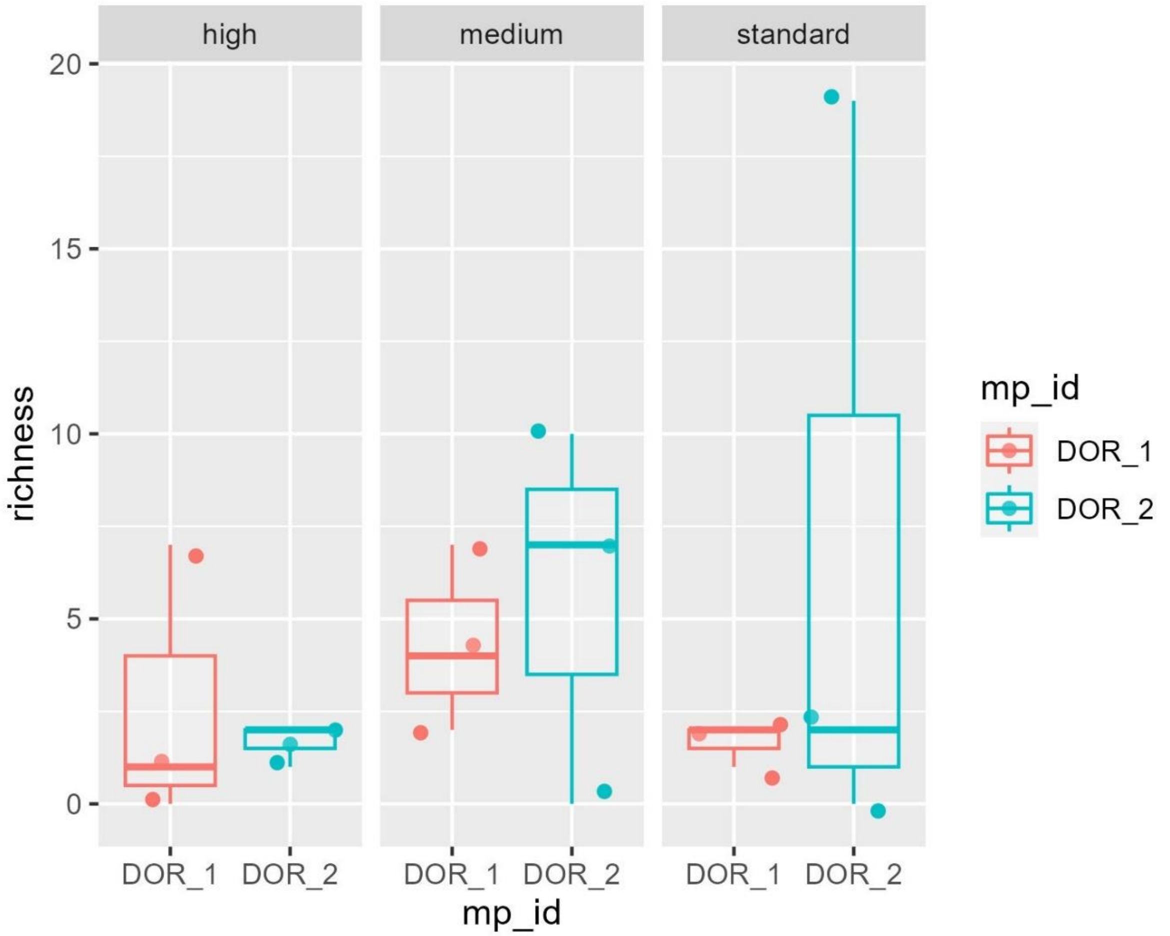

**Fig. S4.**

Box plots of the species richness for samples (DOR_1, DOR_2) collected from Studland Bay, Dorset at the South Beach 1 dive site, sorted by gauze weight ranges.

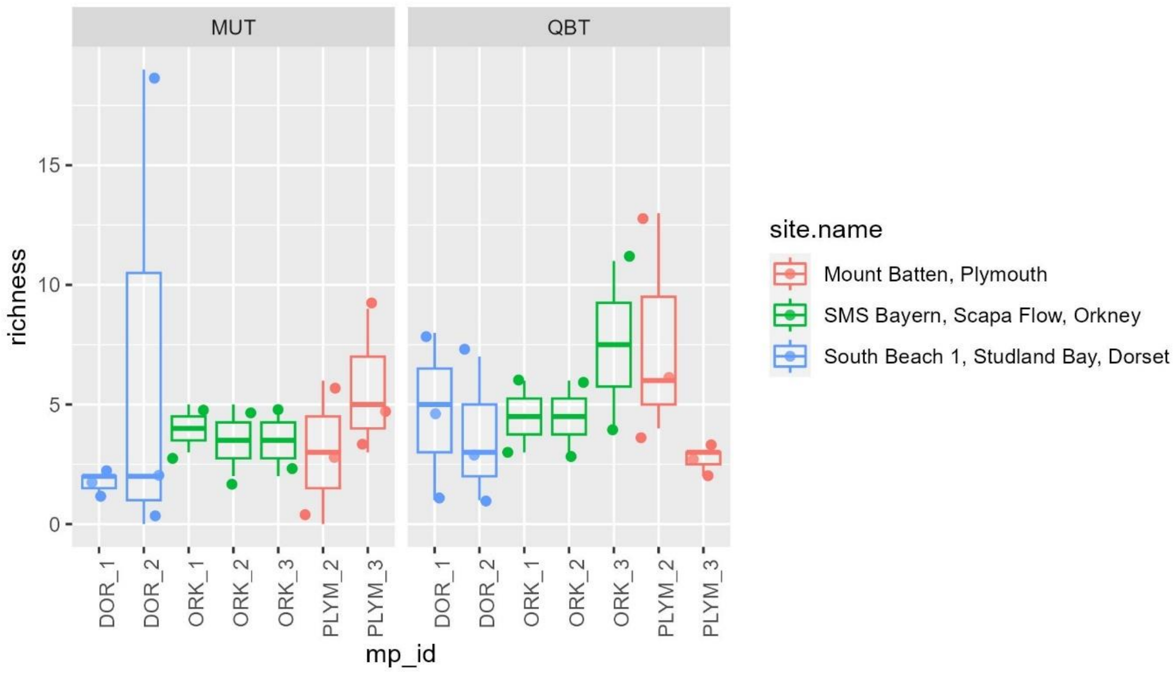

**Fig. S5.**

Species richness for samples collected from Mount Batten, Plymouth, SMS Bayern, Orkney, and South Beach 1, Studland Bay. The Mu-DNA Tissue extraction method (MUT, left) is compared to the Qiagen Blood and Tissue kit (QBT, right).

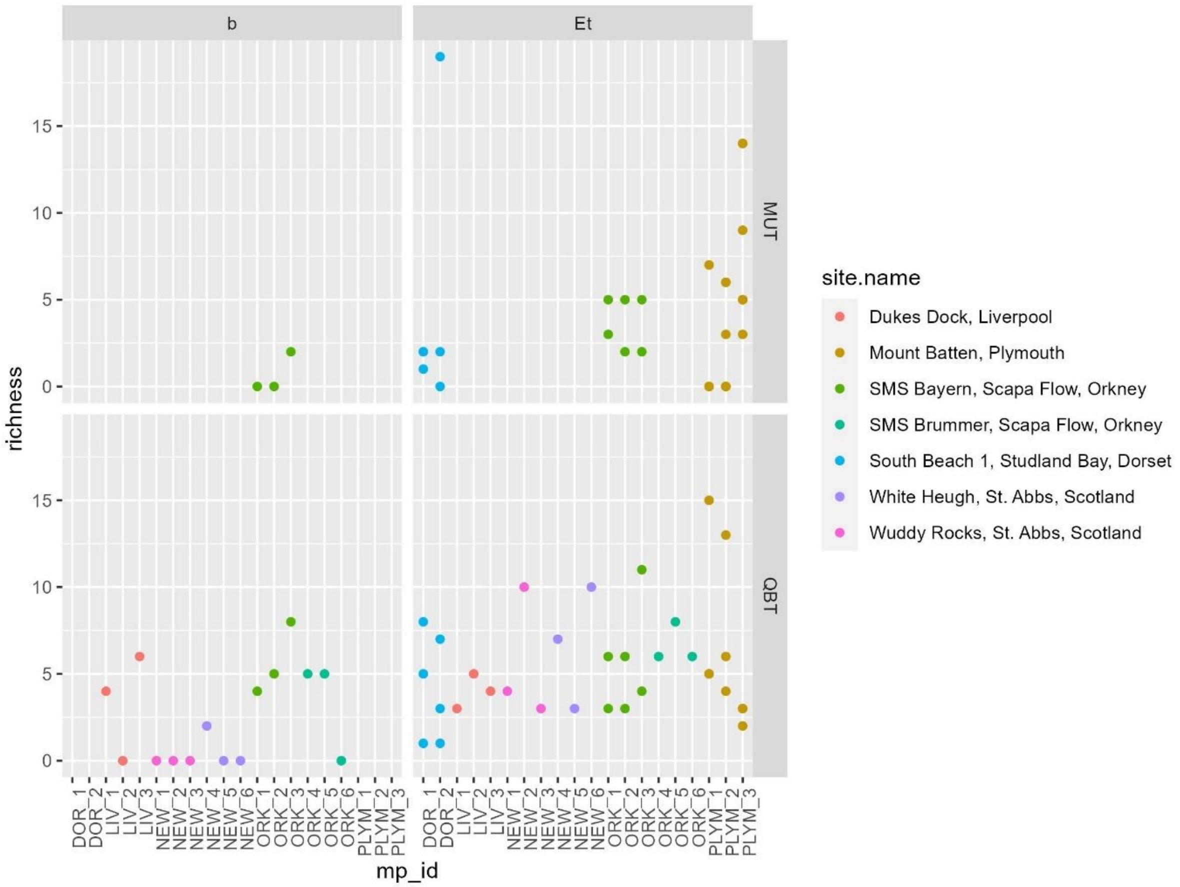

**Fig. S6.**

Stacked bar charts comparing the detected species richness for samples from various dive sites which underwent different preservation treatments (vertical columns), where ‘b’ = silica beads and ‘Et’ = ethanol, and different DNA extraction methods, where ‘MUT’ = Mu-DNA Tissue method and ‘QBT’ = Qiagen Blood and Tissue kit. The horizontal QBT panel shows samples used to test for the effects of the different preservation techniques (Supplementary Table 13). Samples that did not contain detections are shown with a species richness of zero, therefore treatment iterations with no data do not exist.

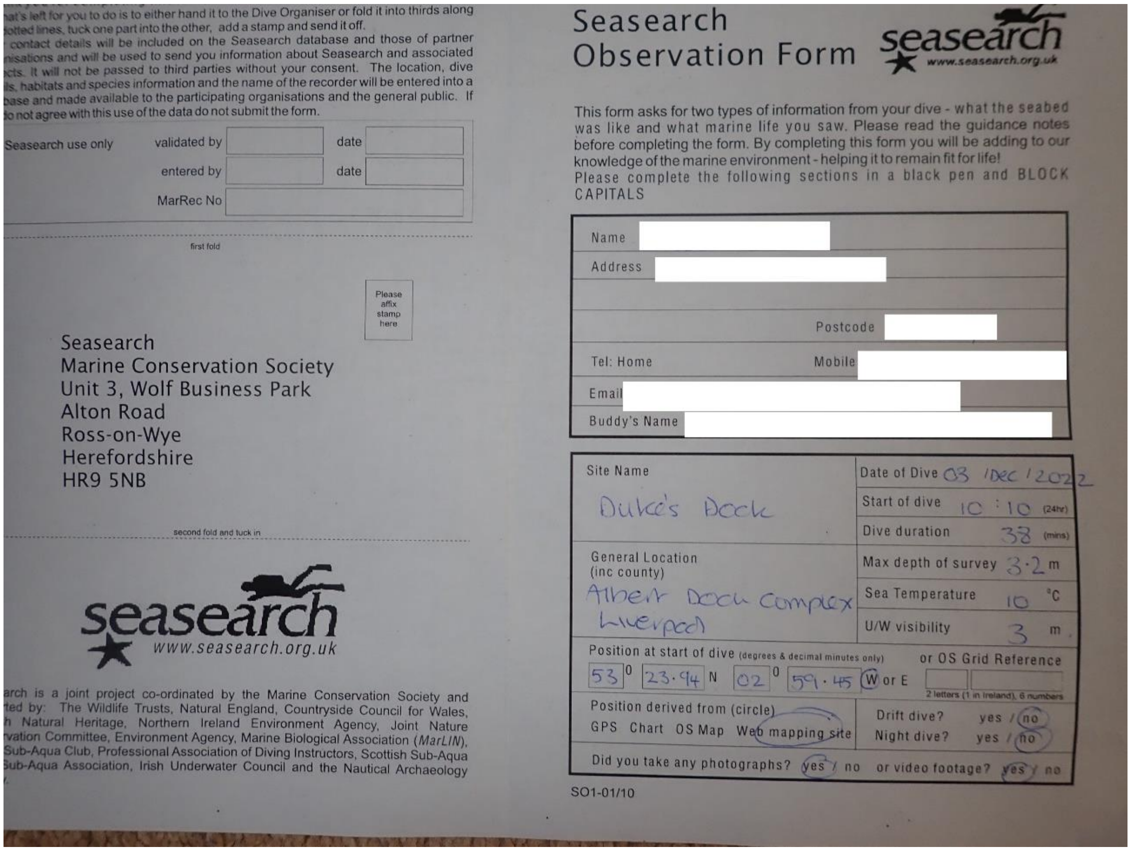
...at's left for you to do is to either hand it to the Dive Organiser or fold it into thirds along ...otted lines, tuck one part into the other, add a stamp and send it off.
... contact details will be included on the Seasearch database and those of partner ...nisations and will be used to send you information about Seasearch and associated ...cts. It will not be passed to third parties without your consent. The location, dive ...ils, habitats and species information and the name of the recorder will be entered into a ...base and made available to the participating organisations and the general public. If ...do not agree with this use of the data do not submit the form.

| Seasearch use only | validated by | | date |
|---|---|---|---|
| | entered by | | date |
| | MarRec No | | |

first fold

Please affix stamp here

Seasearch
Marine Conservation Society
Unit 3, Wolf Business Park
Alton Road
Ross-on-Wye
Herefordshire
HR9 5NB

second fold and tuck in

seasearch
www.seasearch.org.uk

...arch is a joint project co-ordinated by the Marine Conservation Society and ...ted by: The Wildlife Trusts, Natural England, Countryside Council for Wales, ...h Natural Heritage, Northern Ireland Environment Agency, Joint Nature ...rvation Committee, Environment Agency, Marine Biological Association (*MarLIN*), ...Sub-Aqua Club, Professional Association of Diving Instructors, Scottish Sub-Aqua ...Sub-Aqua Association, Irish Underwater Council and the Nautical Archaeology ...

### Seasearch Observation Form

seasearch
www.seasearch.org.uk

This form asks for two types of information from your dive - what the seabed was like and what marine life you saw. Please read the guidance notes before completing the form. By completing this form you will be adding to our knowledge of the marine environment - helping it to remain fit for life!
Please complete the following sections in a black pen and BLOCK CAPITALS

Name
Address
Postcode
Tel: Home Mobile
Email
Buddy's Name

| Site Name | Date of Dive 03 / Dec / 2022 |
|---|---|
| Duke's Dock | Start of dive 10 : 10 (24hr) |
| | Dive duration 38 (mins) |
| General Location (inc county) | Max depth of survey 3.2 m |
| Albert Dock Complex Liverpool | Sea Temperature 10 °C |
| | U/W visibility 3 m |

Position at start of dive (degrees & decimal minutes only): 53° 23.94 N 02° 59.45 W or E — or OS Grid Reference (2 letters (1 in Ireland), 6 numbers)

Position derived from (circle): GPS Chart OS Map Web mapping site
Drift dive? yes / no
Night dive? yes / no
Did you take any photographs? yes / no — or video footage? yes / no

SO1-01/10

**Fig. S7.**

Seasearch Obervation form from the Dukes Dock, Liverpool dive (December 2022). Front side shown above and back side on the following page.

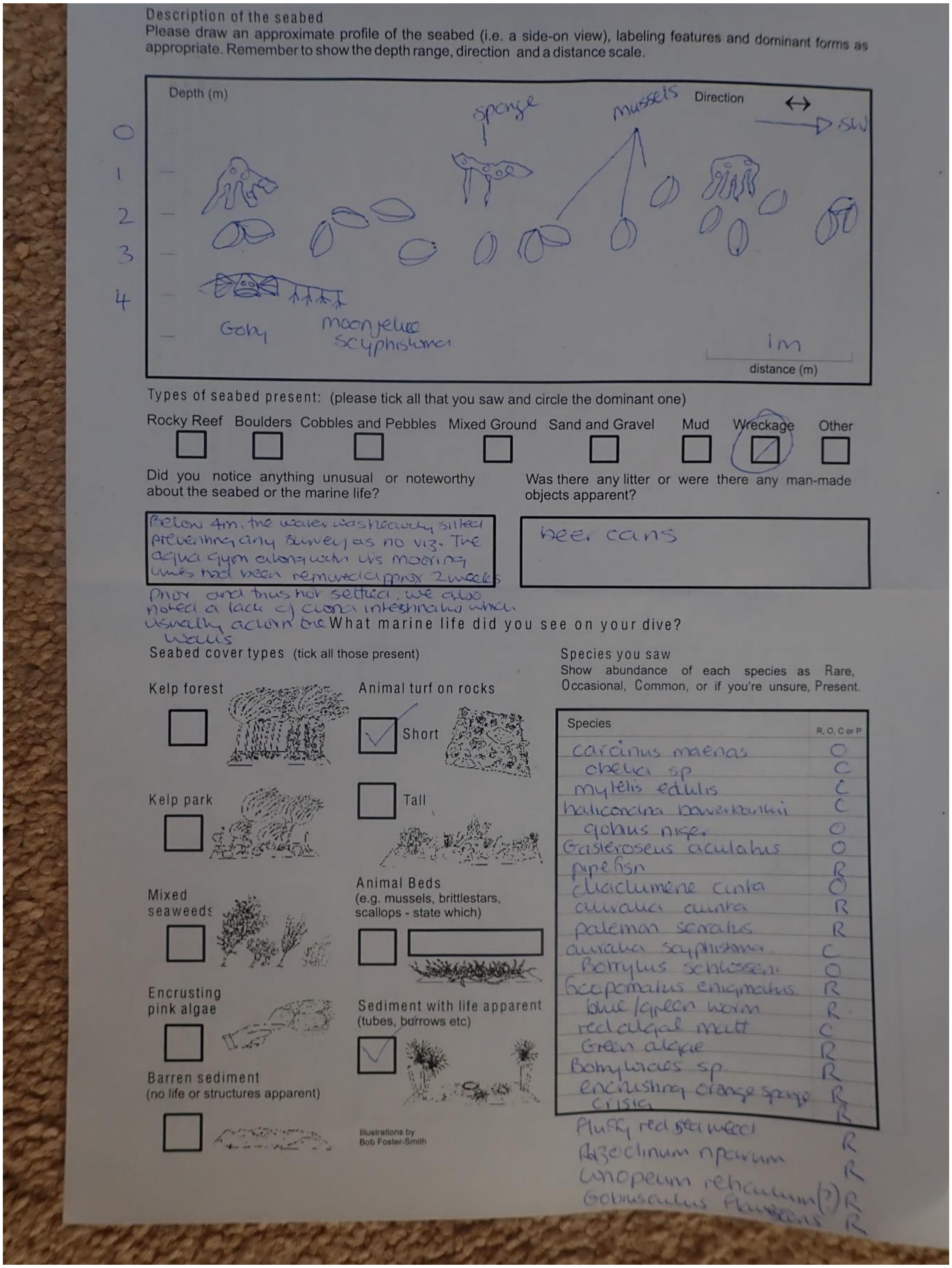

Description of the seabed

Please draw an approximate profile of the seabed (i.e. a side-on view), labeling features and dominant forms as appropriate. Remember to show the depth range, direction and a distance scale.

Types of seabed present: (please tick all that you saw and circle the dominant one)

Rocky Reef ☐ Boulders ☐ Cobbles and Pebbles ☐ Mixed Ground ☐ Sand and Gravel ☐ Mud ☐ Wreckage ☑ (circled) Other ☐

Did you notice anything unusual or noteworthy about the seabed or the marine life?

Below 4m, the water was heavily silted preventing any survey as no viz. The aqua gym along with its mooring lines had been removed approx 2 weeks prior and thus not settled. We also noted a lack of ciona intestinalis which usually adorn the walls

Was there any litter or were there any man-made objects apparent?

beer cans

What marine life did you see on your dive?

Seabed cover types (tick all those present)

Kelp forest ☐

Kelp park ☐

Mixed seaweeds ☐

Encrusting pink algae ☐

Barren sediment (no life or structures apparent) ☐

Animal turf on rocks: Short ☑ Tall ☐

Animal Beds (e.g. mussels, brittlestars, scallops - state which) ☐

Sediment with life apparent (tubes, burrows etc) ☑

Illustrations by Bob Foster-Smith

Species you saw

Show abundance of each species as Rare, Occasional, Common, or if you're unsure, Present.

| Species | R, O, C or P |
| --- | --- |
| carcinus maenas | O |
| obelia sp | C |
| mytilis edulis | C |
| halicondria bowerbankii | C |
| gobius niger | O |
| Gasterosteus aculatus | O |
| pipefish | R |
| diadumene cincta | O |
| aurelia aurita | R |
| paleamon serratus | R |
| aurelia scyphistoma | C |
| Botryllus schlosseri | O |
| ficopomatus enigmaticus | R |
| blue/green worm | R |
| red algal mat | C |
| Green algae | R |
| Botrylloides sp | R |
| encrusting orange sponge | R |
| crisia | R |
| Fluffy red sea weed | R |
| Aiptasia/clinum npeum [illegible] | R |
| Lomentaria [illegible] (?) | R |
| Gobiusculus flavescens | R |

**Fig. S7 (continued).**

Seasearch Obervation form from the Dukes Dock, Liverpool dive (December 2022).

**Table S1.**

Sequenced samples and metadata used for analysis.

**S1.1** Sample information for all 190 samples sequenced where 'Num' = sample number, 'Run' = sequencing run, 'Type' = sample type ('MP' = metaprobe, 'eDNA' = eDNA filter), 'mp_id' = metaprobe ID, 'Pres' = preservation treatment ('Et' = 100% ethanol, 'b' = silica beads), 'Ext' = extraction type, 'Gau' = gauze treatment, 'W' = weight in grams, and 'ICES area' = International Council for Exploration of the Sea advisory areas. The abbreviations in the 'Gau' column indicate the different ways gauze had been rolled and placed in the metaprobe: 'ham' = cutting rolls in thirds by width (3.3 cm x 3.7 m); 'hotf' = cutting rolls in half by length and width (5 cm x 1.85 m); 'hott' = cutting rolls in thirds by length (10 cm x 1.23 m); 'whole' = no modification (10 cm x 3.7 m).

| Num | Run | Type | mp_id | Dive Site Name | Location | Country | Via | Pres | Ext | Gau | W (g) | Weight Category | Primer | Time (min) | ICES area |
|---|---|---|---|---|---|---|---|---|---|---|---|---|---|---|---|
| 1 | A | MP | PLYM_1 | Mount Batten | Plymouth | England | snorkel | Et | QBT | ham | 0.25 | standard | tele02 | 60 | 7.e |
| 2 | A | MP | PLYM_1 | Mount Batten | Plymouth | England | snorkel | Et | QBT | ham | 0.2 | standard | tele02 | 60 | 7.e |
| 3 | A | MP | PLYM_1 | Mount Batten | Plymouth | England | snorkel | Et | QBT | ham | 0.4 | standard | tele02 | 60 | 7.e |
| 4 | A | MP | PLYM_2 | Mount Batten | Plymouth | England | snorkel | Et | QBT | ham | 0.31 | standard | tele02 | 60 | 7.e |
| 5 | A | MP | PLYM_2 | Mount Batten | Plymouth | England | snorkel | Et | QBT | ham | 0.27 | standard | tele02 | 60 | 7.e |
| 6 | A | MP | PLYM_2 | Mount Batten | Plymouth | England | snorkel | Et | QBT | ham | 0.31 | standard | tele02 | 60 | 7.e |
| 7 | A | MP | PLYM_3 | Mount Batten | Plymouth | England | snorkel | Et | QBT | ham | 0.21 | standard | tele02 | 60 | 7.e |
| 8 | A | MP | PLYM_3 | Mount Batten | Plymouth | England | snorkel | Et | QBT | ham | 0.22 | standard | tele02 | 60 | 7.e |
| 9 | A | MP | PLYM_3 | Mount Batten | Plymouth | England | snorkel | Et | QBT | ham | 0.36 | standard | tele02 | 60 | 7.e |
| 10 | A | MP | DOR_1 | South Beach 1, Studland Bay | Dorset | England | SCUBA | Et | QBT | hotf | 0.2 | standard | tele02 | 77 | 7.d |
| 11 | A | MP | DOR_1 | South Beach 1, Studland Bay | Dorset | England | SCUBA | Et | QBT | hotf | 0.26 | standard | tele02 | 77 | 7.d |
| 12 | A | MP | DOR_1 | South Beach 1, Studland Bay | Dorset | England | SCUBA | Et | QBT | hotf | 0.2 | standard | tele02 | 77 | 7.d |
| 13 | A | MP | DOR_2 | South Beach 1, Studland Bay | Dorset | England | SCUBA | Et | QBT | hotf | 0.28 | standard | tele02 | 90 | 7.d |
| 14 | A | MP | DOR_2 | South Beach 1, Studland Bay | Dorset | England | SCUBA | Et | QBT | hotf | 0.4 | standard | tele02 | 90 | 7.d |
| 15 | A | MP | DOR_2 | South Beach 1, Studland Bay | Dorset | England | SCUBA | Et | QBT | hotf | 0.21 | standard | tele02 | 90 | 7.d |

| | | | | | | | | | | | | | | | |
|---|---|---|---|---|---|---|---|---|---|---|---|---|---|---|---|
| 16 | A | MP | DOR_3 | South Beach 2, Studland Bay | Dorset | England | SCUBA | Et | QBT | hotf | 0.38 | standard | tele02 | 73 | 7.d |
| 17 | A | MP | DOR_3 | South Beach 2, Studland Bay | Dorset | England | SCUBA | Et | QBT | hotf | 0.22 | standard | tele02 | 73 | 7.d |
| 18 | A | MP | DOR_3 | South Beach 2, Studland Bay | Dorset | England | SCUBA | Et | QBT | hotf | 0.23 | standard | tele02 | 73 | 7.d |
| 19 | A | MP | DOR_4 | South Beach 2, Studland Bay | Dorset | England | SCUBA | Et | QBT | hotf | 0.38 | standard | tele02 | 73 | 7.d |
| 20 | A | MP | DOR_4 | South Beach 2, Studland Bay | Dorset | England | SCUBA | Et | QBT | hotf | 0.2 | standard | tele02 | 73 | 7.d |
| 21 | A | MP | DOR_4 | South Beach 2, Studland Bay | Dorset | England | SCUBA | Et | QBT | hotf | 0.4 | standard | tele02 | 73 | 7.d |
| 22 | A | MP | PLYM_2 | Mount Batten | Plymouth | England | snorkel | Et | MUT | ham | NR | standard | tele02 | 60 | 7.e |
| 23 | A | MP | PLYM_2 | Mount Batten | Plymouth | England | snorkel | Et | MUT | ham | NR | standard | tele02 | 60 | 7.e |
| 24 | A | MP | PLYM_2 | Mount Batten | Plymouth | England | snorkel | Et | MUT | ham | NR | standard | tele02 | 60 | 7.e |
| 25 | A | MP | PLYM_3 | Mount Batten | Plymouth | England | snorkel | Et | MUT | ham | NR | standard | tele02 | 60 | 7.e |
| 26 | A | MP | PLYM_3 | Mount Batten | Plymouth | England | snorkel | Et | MUT | ham | NR | standard | tele02 | 60 | 7.e |
| 27 | A | MP | PLYM_3 | Mount Batten | Plymouth | England | snorkel | Et | MUT | ham | NR | standard | tele02 | 60 | 7.e |
| 28 | A | MP | DOR_1 | South Beach 1, Studland Bay | Dorset | England | SCUBA | Et | MUT | hotf | 0.31 | standard | tele02 | 77 | 7.d |
| 29 | A | MP | DOR_1 | South Beach 1, Studland Bay | Dorset | England | SCUBA | Et | MUT | hotf | 0.21 | standard | tele02 | 77 | 7.d |
| 30 | A | MP | DOR_1 | South Beach 1, Studland Bay | Dorset | England | SCUBA | Et | MUT | hotf | 0.2 | standard | tele02 | 77 | 7.d |
| 31 | A | MP | DOR_2 | South Beach 1, Studland Bay | Dorset | England | SCUBA | Et | MUT | hotf | 0.24 | standard | tele02 | 90 | 7.d |
| 32 | A | MP | DOR_2 | South Beach 1, Studland Bay | Dorset | England | SCUBA | Et | MUT | hotf | 0.24 | standard | tele02 | 90 | 7.d |
| 33 | A | MP | DOR_2 | South Beach 1, Studland Bay | Dorset | England | SCUBA | Et | MUT | hotf | 0.32 | standard | tele02 | 90 | 7.d |
| 34 | A | MP | DOR_1 | South Beach 1, Studland Bay | Dorset | England | SCUBA | Et | MUT | hotf | 0.79 | medium | tele02 | 77 | 7.d |
| 35 | A | MP | DOR_1 | South Beach 1, Studland Bay | Dorset | England | SCUBA | Et | MUT | hotf | 0.61 | medium | tele02 | 77 | 7.d |
| 36 | A | MP | DOR_1 | South Beach 1, Studland Bay | Dorset | England | SCUBA | Et | MUT | hotf | 0.78 | medium | tele02 | 77 | 7.d |

| | | | | | | | | | | | | | | | |
|---|---|---|---|---|---|---|---|---|---|---|---|---|---|---|---|
| 37 | A | MP | DOR_2 | South Beach 1, Studland Bay | Dorset | England | SCUBA | Et | MUT | hotf | 0.61 | medium | tele02 | 90 | 7.d |
| 38 | A | MP | DOR_2 | South Beach 1, Studland Bay | Dorset | England | SCUBA | Et | MUT | hotf | 0.64 | medium | tele02 | 90 | 7.d |
| 39 | A | MP | DOR_2 | South Beach 1, Studland Bay | Dorset | England | SCUBA | Et | MUT | hotf | 0.7 | medium | tele02 | 90 | 7.d |
| 40 | A | MP | DOR_1 | South Beach 1, Studland Bay | Dorset | England | SCUBA | Et | MUT | hotf | 1.05 | high | tele02 | 77 | 7.d |
| 41 | A | MP | DOR_1 | South Beach 1, Studland Bay | Dorset | England | SCUBA | Et | MUT | hotf | 1.03 | high | tele02 | 77 | 7.d |
| 42 | A | MP | DOR_1 | South Beach 1, Studland Bay | Dorset | England | SCUBA | Et | MUT | hotf | 1 | high | tele02 | 77 | 7.d |
| 43 | A | MP | DOR_2 | South Beach 1, Studland Bay | Dorset | England | SCUBA | Et | MUT | hotf | 1.04 | high | tele02 | 90 | 7.d |
| 44 | A | MP | DOR_2 | South Beach 1, Studland Bay | Dorset | England | SCUBA | Et | MUT | hotf | 1.1 | high | tele02 | 90 | 7.d |
| 45 | A | MP | DOR_2 | South Beach 1, Studland Bay | Dorset | England | SCUBA | Et | MUT | hotf | 1.01 | high | tele02 | 90 | 7.d |
| 46 | A | eDNA | NA | SMS Bayern, Scapa Flow | Orkney | Scotland | NA | NA | QBT | NA | NA | NA | tele02 | NA | 4.a |
| 47 | A | eDNA | NA | SMS Bayern, Scapa Flow | Orkney | Scotland | NA | NA | QBT | NA | NA | NA | tele02 | NA | 4.a |
| 48 | A | eDNA | NA | SMS Bayern, Scapa Flow | Orkney | Scotland | NA | NA | QBT | NA | NA | NA | tele02 | NA | 4.a |
| 49 | A | eDNA | NA | SMS Bayern, Scapa Flow | Orkney | Scotland | NA | NA | QBT | NA | NA | NA | tele02 | NA | 4.a |
| 50 | A | MP | ORK_1 | SMS Bayern, Scapa Flow | Orkney | Scotland | SCUBA | Et | QBT | hott | 0.23 | standard | tele02 | NR | 4.a |
| 51 | A | MP | ORK_1 | SMS Bayern, Scapa Flow | Orkney | Scotland | SCUBA | Et | MUT | hott | 0.2 | standard | tele02 | NR | 4.a |
| 52 | A | MP | ORK_1 | SMS Bayern, Scapa Flow | Orkney | Scotland | SCUBA | Et | QBT | hott | 0.27 | standard | tele02 | NR | 4.a |
| 53 | A | MP | ORK_1 | SMS Bayern, Scapa Flow | Orkney | Scotland | SCUBA | Et | MUT | hott | 0.2 | standard | tele02 | NR | 4.a |
| 54 | A | MP | ORK_1 | SMS Bayern, Scapa Flow | Orkney | Scotland | SCUBA | b | QBT | hott | 0.08 | standard | tele02 | NR | 4.a |
| 55 | A | MP | ORK_1 | SMS Bayern, Scapa Flow | Orkney | Scotland | SCUBA | b | MUT | hott | 0.06 | standard | tele02 | NR | 4.a |

| | | | | | | | | | | | | | | | |
|---|---|---|---|---|---|---|---|---|---|---|---|---|---|---|---|
| 56 | A | MP | ORK_2 | SMS Bayern, Scapa Flow | Orkney | Scotland | SCUBA | Et | QBT | hott | 0.27 | standard | tele02 | NR | 4.a |
| 57 | A | MP | ORK_2 | SMS Bayern, Scapa Flow | Orkney | Scotland | SCUBA | Et | MUT | hott | 0.23 | standard | tele02 | NR | 4.a |
| 58 | A | MP | ORK_2 | SMS Bayern, Scapa Flow | Orkney | Scotland | SCUBA | Et | QBT | hott | 0.32 | standard | tele02 | NR | 4.a |
| 59 | A | MP | ORK_2 | SMS Bayern, Scapa Flow | Orkney | Scotland | SCUBA | Et | MUT | hott | 0.29 | standard | tele02 | NR | 4.a |
| 60 | A | MP | ORK_2 | SMS Bayern, Scapa Flow | Orkney | Scotland | SCUBA | b | QBT | hott | 0.08 | standard | tele02 | NR | 4.a |
| 61 | A | MP | ORK_2 | SMS Bayern, Scapa Flow | Orkney | Scotland | SCUBA | b | MUT | hott | 0.08 | standard | tele02 | NR | 4.a |
| 62 | A | MP | ORK_3 | SMS Bayern, Scapa Flow | Orkney | Scotland | SCUBA | Et | QBT | hott | 0.2 | standard | tele02 | NR | 4.a |
| 63 | A | MP | ORK_3 | SMS Bayern, Scapa Flow | Orkney | Scotland | SCUBA | Et | MUT | hott | 0.2 | standard | tele02 | NR | 4.a |
| 64 | A | MP | ORK_3 | SMS Bayern, Scapa Flow | Orkney | Scotland | SCUBA | Et | QBT | hott | 0.36 | standard | tele02 | NR | 4.a |
| 65 | A | MP | ORK_3 | SMS Bayern, Scapa Flow | Orkney | Scotland | SCUBA | Et | MUT | hott | 0.26 | standard | tele02 | NR | 4.a |
| 66 | A | MP | ORK_3 | SMS Bayern, Scapa Flow | Orkney | Scotland | SCUBA | b | QBT | hott | 0.15 | standard | tele02 | NR | 4.a |
| 67 | A | MP | ORK_3 | SMS Bayern, Scapa Flow | Orkney | Scotland | SCUBA | b | MUT | hott | 0.13 | standard | tele02 | NR | 4.a |
| 68 | B | MP | BPD_7 | Ocean Exhibit, Blue Planet, Ellesmere Port | Blue Planet Aquarium | shark tank | soak | Et | QBT | whole | NR | standard | elas02 | 60 | NA |
| 69 | B | MP | BPD_8 | Ocean Exhibit, Blue Planet, Ellesmere Port | Blue Planet Aquarium | shark tank | soak | Et | QBT | whole | NR | standard | elas02 | 120 | NA |
| 70 | B | MP | BPD_8 | Ocean Exhibit, Blue Planet, Ellesmere Port | Blue Planet Aquarium | shark tank | soak | Et | QBT | whole | NR | standard | elas02 | 120 | NA |
| 71 | B | MP | BPD_9 | Ocean Exhibit, Blue Planet, Ellesmere Port | Blue Planet Aquarium | shark tank | soak | Et | QBT | whole | NR | standard | elas02 | 240 | NA |

| | | | | | | | | | | | | | | | |
|---|---|---|---|---|---|---|---|---|---|---|---|---|---|---|---|
| 72 | B | MP | BPD_9 | Ocean Exhibit, Blue Planet, Ellesmere Port | Blue Planet Aquarium | shark tank | soak | Et | QBT | whole | NR | standard | elas02 | 240 | NA |
| 73 | B | MP | BPD_1 | Ocean Exhibit, Blue Planet, Ellesmere Port | Blue Planet Aquarium | dive2 | SCUBA | Et | QBT | whole | NR | standard | elas02 | 65 | NA |
| 74 | B | MP | BPD_1 | Ocean Exhibit, Blue Planet, Ellesmere Port | Blue Planet Aquarium | dive2 | SCUBA | Et | QBT | whole | NR | standard | elas02 | 65 | NA |
| 75 | B | MP | BPD_2 | Ocean Exhibit, Blue Planet, Ellesmere Port | Blue Planet Aquarium | dive2 | SCUBA | Et | QBT | whole | NR | standard | elas02 | 65 | NA |
| 76 | B | MP | BPD_2 | Ocean Exhibit, Blue Planet, Ellesmere Port | Blue Planet Aquarium | dive2 | SCUBA | Et | QBT | whole | NR | standard | elas02 | 65 | NA |
| 77 | B | eDNA | NA | Ocean Exhibit, Blue Planet, Ellesmere Port | Blue Planet Aquarium | shark tank | NA | NA | QBT | NA | NA | NA | elas02 | NA | NA |
| 78 | B | eDNA | NA | Ocean Exhibit, Blue Planet, Ellesmere Port | Blue Planet Aquarium | shark tank | NA | NA | QBT | NA | NA | NA | elas02 | NA | NA |
| 79 | B | eDNA | NA | Ocean Exhibit, Blue Planet, Ellesmere Port | Blue Planet Aquarium | shark tank | NA | NA | QBT | NA | NA | NA | elas02 | NA | NA |
| 80 | B | eDNA | NA | Ocean Exhibit, Blue Planet, Ellesmere Port | Blue Planet Aquarium | shark tank | NA | NA | QBT | NA | NA | NA | elas02 | NA | NA |
| 81 | B | MP | BPD_3 | Ocean Exhibit, Blue Planet, Ellesmere Port | Blue Planet Aquarium | dive1 | SCUBA | Et | QBT | whole | NR | standard | elas02 | 50 | NA |
| 82 | B | MP | BPD_3 | Ocean Exhibit, Blue Planet, Ellesmere Port | Blue Planet Aquarium | dive1 | SCUBA | Et | QBT | whole | NR | standard | elas02 | 50 | NA |
| 83 | B | MP | BPD_4 | Ocean Exhibit, Blue Planet, Ellesmere Port | Blue Planet Aquarium | dive1 | SCUBA | Et | QBT | whole | NR | standard | elas02 | 50 | NA |

| | | | | | | | | | | | | | | | |
|---|---|---|---|---|---|---|---|---|---|---|---|---|---|---|---|
| 84 | B | MP | BPD_4 | Ocean Exhibit, Blue Planet, Ellesmere Port | Blue Planet Aquarium | dive1 | SCUBA | Et | QBT | whole | NR | standard | elas02 | 50 | NA |
| 85 | B | MP | BPD_5 | Ocean Exhibit, Blue Planet, Ellesmere Port | Blue Planet Aquarium | shark tank | soak | Et | QBT | whole | NR | standard | elas02 | 10 | NA |
| 86 | B | MP | BPD_5 | Ocean Exhibit, Blue Planet, Ellesmere Port | Blue Planet Aquarium | shark tank | soak | Et | QBT | whole | NR | standard | elas02 | 10 | NA |
| 87 | B | MP | BPD_6 | Ocean Exhibit, Blue Planet, Ellesmere Port | Blue Planet Aquarium | shark tank | soak | Et | QBT | whole | NR | standard | elas02 | 30 | NA |
| 88 | B | MP | BPD_6 | Ocean Exhibit, Blue Planet, Ellesmere Port | Blue Planet Aquarium | shark tank | soak | Et | QBT | whole | NR | standard | elas02 | 30 | NA |
| 89 | B | MP | BPD_7 | Ocean Exhibit, Blue Planet, Ellesmere Port | Blue Planet Aquarium | shark tank | soak | Et | QBT | whole | NR | standard | elas02 | 60 | NA |
| 90 | B | eDNA | NA | SMS Brummer, Scapa Flow | Orkney | Scotland | NA | NA | QBT | NA | NA | NA | elas02 | NA | 4.a |
| 91 | B | eDNA | NA | SMS Brummer, Scapa Flow | Orkney | Scotland | NA | NA | QBT | NA | NA | NA | elas02 | NA | 4.a |
| 92 | B | eDNA | NA | SMS Brummer, Scapa Flow | Orkney | Scotland | NA | NA | QBT | NA | NA | NA | elas02 | NA | 4.a |
| 93 | B | eDNA | NA | SMS Brummer, Scapa Flow | Orkney | Scotland | NA | NA | QBT | NA | NA | NA | elas02 | NA | 4.a |
| 94 | B | MP | ORK_4 | SMS Brummer, Scapa Flow | Orkney | Scotland | SCUBA | Et | QBT | hott | NR | standard | elas02 | 32 | 4.a |
| 95 | B | MP | ORK_5 | SMS Brummer, Scapa Flow | Orkney | Scotland | SCUBA | Et | QBT | hott | NR | standard | elas02 | 32 | 4.a |
| 96 | B | MP | ORK_6 | SMS Brummer, Scapa Flow | Orkney | Scotland | SCUBA | Et | QBT | hott | NR | standard | elas02 | 32 | 4.a |
| 97 | B | MP | ORK_5 | SMS Brummer, Scapa Flow | Orkney | Scotland | SCUBA | b | QBT | hott | NR | standard | elas02 | 32 | 4.a |
| 98 | B | MP | ORK_6 | SMS Brummer, Scapa Flow | Orkney | Scotland | SCUBA | b | QBT | hott | NR | standard | elas02 | 32 | 4.a |
| 99 | B | MP | ORK_4 | SMS Brummer, Scapa Flow | Orkney | Scotland | SCUBA | b | QBT | hott | NR | standard | elas02 | 32 | 4.a |

| | | | | | | | | | | | | | | | |
|---|---|---|---|---|---|---|---|---|---|---|---|---|---|---|---|
| 100 | B | MP | BPC_1 | Coral Cave, Blue Planet, Ellesmere Port | Blue Planet Aquarium | coral cave | soak | Et | QBT | whole | NR | standard | tele02 | 10 | NA |
| 101 | B | MP | BPC_1 | Coral Cave, Blue Planet, Ellesmere Port | Blue Planet Aquarium | coral cave | soak | Et | QBT | whole | NR | standard | tele02 | 10 | NA |
| 102 | B | MP | BPC_2 | Coral Cave, Blue Planet, Ellesmere Port | Blue Planet Aquarium | coral cave | soak | Et | QBT | whole | NR | standard | tele02 | 30 | NA |
| 103 | B | MP | BPC_2 | Coral Cave, Blue Planet, Ellesmere Port | Blue Planet Aquarium | coral cave | soak | Et | QBT | whole | NR | standard | tele02 | 30 | NA |
| 104 | B | MP | BPC_3 | Coral Cave, Blue Planet, Ellesmere Port | Blue Planet Aquarium | coral cave | soak | Et | QBT | whole | NR | standard | tele02 | 60 | NA |
| 105 | B | MP | BPC_3 | Coral Cave, Blue Planet, Ellesmere Port | Blue Planet Aquarium | coral cave | soak | Et | QBT | whole | NR | standard | tele02 | 60 | NA |
| 106 | B | MP | BPC_4 | Coral Cave, Blue Planet, Ellesmere Port | Blue Planet Aquarium | coral cave | soak | Et | QBT | whole | NR | standard | tele02 | 120 | NA |
| 107 | B | MP | BPC_4 | Coral Cave, Blue Planet, Ellesmere Port | Blue Planet Aquarium | coral cave | soak | Et | QBT | whole | NR | standard | tele02 | 120 | NA |
| 108 | B | MP | BPC_5 | Coral Cave, Blue Planet, Ellesmere Port | Blue Planet Aquarium | coral cave | soak | Et | QBT | whole | NR | standard | tele02 | 240 | NA |
| 109 | B | MP | BPC_5 | Coral Cave, Blue Planet, Ellesmere Port | Blue Planet Aquarium | coral cave | soak | Et | QBT | whole | NR | standard | tele02 | 240 | NA |
| 110 | B | eDNA | NA | Dukes Dock | Liverpool | England | NA | NA | QBT | NA | NA | NA | tele02 | NA | 7.a |
| 111 | B | eDNA | NA | Dukes Dock | Liverpool | England | NA | NA | QBT | NA | NA | NA | tele02 | NA | 7.a |
| 112 | B | eDNA | NA | Dukes Dock | Liverpool | England | NA | NA | QBT | NA | NA | NA | tele02 | NA | 7.a |
| 113 | B | eDNA | NA | Dukes Dock | Liverpool | England | NA | NA | QBT | NA | NA | NA | tele02 | NA | 7.a |
| 114 | B | MP | LIV_1 | Dukes Dock | Liverpool | England | SCUBA | Et | QBT | hott | NR | standard | tele02 | 38 | 7.a |
| 115 | B | MP | LIV_2 | Dukes Dock | Liverpool | England | SCUBA | Et | QBT | hott | NR | standard | tele02 | 38 | 7.a |

| | | | | | | | | | | | | | | | |
|---|---|---|---|---|---|---|---|---|---|---|---|---|---|---|---|
| 116 | B | MP | LIV_3 | Dukes Dock | Liverpool | England | SCUBA | Et | QBT | hott | NR | standard | tele02 | 38 | 7.a |
| 117 | B | MP | LIV_3 | Dukes Dock | Liverpool | England | SCUBA | b | QBT | hott | NR | standard | tele02 | 38 | 7.a |
| 118 | B | MP | LIV_2 | Dukes Dock | Liverpool | England | SCUBA | b | QBT | hott | NR | standard | tele02 | 38 | 7.a |
| 119 | B | MP | LIV_1 | Dukes Dock | Liverpool | England | SCUBA | b | QBT | hott | NR | standard | tele02 | 38 | 7.a |
| 120 | B | MP | NEW_1 | Wuddy Rocks | St. Abbs Head | Scotland | SCUBA | Et | QBT | hotf | NR | standard | tele02 | 45 | 4.b |
| 121 | B | MP | NEW_1 | Wuddy Rocks | St. Abbs Head | Scotland | SCUBA | b | QBT | hotf | NR | standard | tele02 | 45 | 4.b |
| 122 | B | MP | NEW_2 | Wuddy Rocks | St. Abbs Head | Scotland | SCUBA | Et | QBT | hotf | NR | standard | tele02 | 45 | 4.b |
| 123 | B | MP | NEW_2 | Wuddy Rocks | St. Abbs Head | Scotland | SCUBA | b | QBT | hotf | NR | standard | tele02 | 45 | 4.b |
| 124 | B | MP | NEW_3 | Wuddy Rocks | St. Abbs Head | Scotland | SCUBA | Et | QBT | hotf | NR | standard | tele02 | 45 | 4.b |
| 125 | B | MP | NEW_3 | Wuddy Rocks | St. Abbs Head | Scotland | SCUBA | b | QBT | hotf | NR | standard | tele02 | 45 | 4.b |
| 126 | B | MP | NEW_4 | White Heugh | St. Abbs Head | Scotland | SCUBA | Et | QBT | hotf | NR | standard | tele02 | 37 | 4.b |
| 127 | B | MP | NEW_4 | White Heugh | St. Abbs Head | Scotland | SCUBA | b | QBT | hotf | NR | standard | tele02 | 37 | 4.b |
| 128 | B | MP | NEW_5 | White Heugh | St. Abbs Head | Scotland | SCUBA | Et | QBT | hotf | NR | standard | tele02 | 37 | 4.b |
| 129 | B | MP | NEW_5 | White Heugh | St. Abbs Head | Scotland | SCUBA | b | QBT | hotf | NR | standard | tele02 | 37 | 4.b |
| 130 | B | MP | NEW_6 | White Heugh | St. Abbs Head | Scotland | SCUBA | Et | QBT | hotf | NR | standard | tele02 | 37 | 4.b |
| 131 | B | MP | NEW_6 | White Heugh | St. Abbs Head | Scotland | SCUBA | b | QBT | hotf | NR | standard | tele02 | 37 | 4.b |
| 132 | C | MP | CV_1 | Bodega de choco | Boa Vista | Cape Verde | SCUBA | Et | QBT | hott | NR | standard | elas02 | 40 | NA |
| 133 | C | MP | CV_1 | Bodega de choco | Boa Vista | Cape Verde | SCUBA | Et | QBT | hott | NR | standard | elas02 | 40 | NA |
| 134 | C | MP | CV_1 | Bodega de choco | Boa Vista | Cape Verde | SCUBA | Et | QBT | hott | NR | standard | elas02 | 40 | NA |
| 135 | C | MP | CA_1 | San Carlos Beach Wall | California | USA | SCUBA | Et | QBT | hotf | NR | standard | elas02 | 42 | NA |

| | | | | | | | | | | | | | | | |
|---|---|---|---|---|---|---|---|---|---|---|---|---|---|---|---|
| 136 | C | MP | CA_1 | San Carlos Beach Wall | California | USA | SCUBA | Et | QBT | hotf | NR | standard | elas02 | 42 | NA |
| 137 | C | MP | CA_1 | San Carlos Beach Wall | California | USA | SCUBA | Et | QBT | hotf | NR | standard | elas02 | 42 | NA |
| 138 | C | MP | RS_1 | South Beach | Gulf of Aqaba | Jordan | SCUBA | Et | QBT | hott | NR | standard | elas02 | 50 | NA |
| 139 | C | MP | RS_1 | South Beach | Gulf of Aqaba | Jordan | SCUBA | Et | QBT | hott | NR | standard | elas02 | 50 | NA |
| 140 | C | MP | RS_1 | South Beach | Gulf of Aqaba | Jordan | SCUBA | Et | QBT | hott | NR | standard | elas02 | 50 | NA |
| 141 | C | MP | SAD_1 | SL9F | Sodwana Bay | South Africa | SCUBA | b | QBT | hotf | NR | standard | elas02 | NR | NA |
| 142 | C | MP | SAD_1 | SL9F | Sodwana Bay | South Africa | SCUBA | b | QBT | hotf | NR | standard | elas02 | NR | NA |
| 143 | C | MP | SAD_1 | SL9F | Sodwana Bay | South Africa | SCUBA | b | QBT | hotf | NR | standard | elas02 | NR | NA |
| 144 | C | MP | SAD_2 | SL9M | Sodwana Bay | South Africa | SCUBA | b | QBT | hotf | NR | standard | elas02 | NR | NA |
| 145 | C | MP | SAD_2 | SL9M | Sodwana Bay | South Africa | SCUBA | b | QBT | hotf | NR | standard | elas02 | NR | NA |
| 146 | C | MP | SAD_2 | SL9M | Sodwana Bay | South Africa | SCUBA | b | QBT | hotf | NR | standard | elas02 | NR | NA |
| 147 | C | MP | SAD_3 | SL7F1 | Sodwana Bay | South Africa | SCUBA | b | QBT | hotf | NR | standard | elas02 | NR | NA |
| 148 | C | MP | SAD_3 | SL7F1 | Sodwana Bay | South Africa | SCUBA | b | QBT | hotf | NR | standard | elas02 | NR | NA |
| 149 | C | MP | SAD_3 | SL7F1 | Sodwana Bay | South Africa | SCUBA | b | QBT | hotf | NR | standard | elas02 | NR | NA |
| 150 | C | MP | SAD_4 | SL7F2 | Sodwana Bay | South Africa | SCUBA | b | QBT | hotf | NR | standard | elas02 | NR | NA |
| 151 | C | MP | SAD_4 | SL7F2 | Sodwana Bay | South Africa | SCUBA | b | QBT | hotf | NR | standard | elas02 | NR | NA |
| 152 | C | MP | SAD_4 | SL7F2 | Sodwana Bay | South Africa | SCUBA | b | QBT | hotf | NR | standard | elas02 | NR | NA |
| 153 | D | MP | NOR_1 | Radbod | Ørsta | Norway | SCUBA | b | MUT | hotf | NR | standard | tele02 | 40 | 2.a.2 |
| 154 | D | MP | NOR_1 | Radbod | Ørsta | Norway | SCUBA | b | MUT | hotf | NR | standard | tele02 | 40 | 2.a.2 |
| 155 | D | MP | NOR_1 | Radbod | Ørsta | Norway | SCUBA | b | MUT | hotf | NR | standard | tele02 | 40 | 2.a.2 |

| | | | | | | | | | | | | | | | |
|---|---|---|---|---|---|---|---|---|---|---|---|---|---|---|---|
| 156 | D | MP | NOR_2 | Måløy | Måløy | Norway | SCUBA | b | MUT | hotf | NR | standard | tele02 | 40 | 4.a |
| 157 | D | MP | NOR_2 | Måløy | Måløy | Norway | SCUBA | b | MUT | hotf | NR | standard | tele02 | 40 | 4.a |
| 158 | D | MP | NOR_2 | Måløy | Måløy | Norway | SCUBA | b | MUT | hotf | NR | standard | tele02 | 40 | 4.a |
| 159 | D | MP | NOR_3 | Dogfish Walk | Bergen | Norway | SCUBA | b | MUT | hotf | NR | standard | tele02 | 41 | 4.a |
| 160 | D | MP | NOR_3 | Dogfish Walk | Bergen | Norway | SCUBA | b | MUT | hotf | NR | standard | tele02 | 41 | 4.a |
| 161 | D | MP | NOR_3 | Dogfish Walk | Bergen | Norway | SCUBA | b | MUT | hotf | NR | standard | tele02 | 41 | 4.a |
| 162 | D | MP | NOR_4 | Dogfish Walk | Bergen | Norway | SCUBA | b | MUT | hotf | NR | standard | tele02 | 41 | 4.a |
| 163 | D | MP | PLYM_1 | Mount Batten | Plymouth | England | snorkel | Et | MUT | ham | NR | standard | tele02 | 60 | 7.e |
| 164 | D | MP | PLYM_1 | Mount Batten | Plymouth | England | snorkel | Et | MUT | ham | NR | standard | tele02 | 60 | 7.e |
| 165 | D | MP | PLYM_1 | Mount Batten | Plymouth | England | snorkel | Et | MUT | ham | NR | standard | tele02 | 60 | 7.e |
| 166 | D | MP | PLYM_2 | Mount Batten | Plymouth | England | snorkel | Et | MUT | ham | NR | standard | tele02 | 60 | 7.e |
| 167 | D | MP | PLYM_2 | Mount Batten | Plymouth | England | snorkel | Et | MUT | ham | NR | standard | tele02 | 60 | 7.e |
| 168 | D | MP | PLYM_2 | Mount Batten | Plymouth | England | snorkel | Et | MUT | ham | NR | standard | tele02 | 60 | 7.e |
| 169 | D | MP | PLYM_3 | Mount Batten | Plymouth | England | snorkel | Et | MUT | ham | NR | standard | tele02 | 60 | 7.e |
| 170 | D | MP | PLYM_3 | Mount Batten | Plymouth | England | snorkel | Et | MUT | ham | NR | standard | tele02 | 60 | 7.e |
| 171 | D | MP | PLYM_3 | Mount Batten | Plymouth | England | snorkel | Et | MUT | ham | NR | standard | tele02 | 60 | 7.e |
| 172 | D | MP | PLYM_4 | Mount Batten | Plymouth | England | snorkel | Et | MUT | ham | NR | standard | tele02 | 60 | 7.e |
| 173 | D | MP | PLYM_4 | Mount Batten | Plymouth | England | snorkel | Et | MUT | ham | NR | standard | tele02 | 60 | 7.e |
| 174 | D | MP | PLYM_4 | Mount Batten | Plymouth | England | snorkel | Et | MUT | ham | NR | standard | tele02 | 60 | 7.e |
| 175 | D | MP | PLYM_5 | Mount Batten | Plymouth | England | snorkel | Et | MUT | ham | NR | standard | tele02 | 60 | 7.e |
| 176 | D | MP | PLYM_5 | Mount Batten | Plymouth | England | snorkel | Et | MUT | ham | NR | standard | tele02 | 60 | 7.e |
| 177 | D | MP | PLYM_5 | Mount Batten | Plymouth | England | snorkel | Et | MUT | ham | NR | standard | tele02 | 60 | 7.e |
| 178 | D | MP | PLYM_6 | Mount Batten | Plymouth | England | snorkel | Et | MUT | ham | NR | standard | tele02 | 60 | 7.e |
| 179 | D | MP | PLYM_6 | Mount Batten | Plymouth | England | snorkel | Et | MUT | ham | NR | standard | tele02 | 60 | 7.e |
| 180 | D | MP | PLYM_6 | Mount Batten | Plymouth | England | snorkel | Et | MUT | ham | NR | standard | tele02 | 60 | 7.e |
| 181 | D | MP | PLYM_7 | Mount Batten | Plymouth | England | snorkel | Et | MUT | ham | NR | standard | tele02 | 60 | 7.e |
| 182 | D | MP | PLYM_7 | Mount Batten | Plymouth | England | snorkel | Et | MUT | ham | NR | standard | tele02 | 60 | 7.e |
| 183 | D | MP | PLYM_7 | Mount Batten | Plymouth | England | snorkel | Et | MUT | ham | NR | standard | tele02 | 60 | 7.e |
| 184 | D | MP | NOR_4 | Dogfish Walk | Bergen | Norway | SCUBA | b | MUT | hotf | NR | standard | tele02 | 41 | 4.a |
| 185 | D | MP | NOR_4 | Dogfish Walk | Bergen | Norway | SCUBA | b | MUT | hotf | NR | standard | tele02 | 41 | 4.a |
| 186 | D | MP | NOR_5 | Dogfish Walk | Bergen | Norway | SCUBA | b | MUT | hotf | NR | standard | tele02 | 41 | 4.a |

| | | | | | | | | | | | | | | | |
|---|---|---|---|---|---|---|---|---|---|---|---|---|---|---|---|
| 187 | D | MP | NOR_5 | Dogfish Walk | Bergen | Norway | SCUBA | b | MUT | hotf | NR | standard | tele02 | 41 | 4.a |
| 188 | D | MP | NOR_5 | Dogfish Walk | Bergen | Norway | SCUBA | b | MUT | hotf | NR | standard | tele02 | 41 | 4.a |
| 189 | D | MP | NOR_6 | Welheim | Florø | Norway | SCUBA | b | MUT | hotf | NR | standard | tele02 | 41 | 4.a |
| 190 | D | MP | NOR_6 | Welheim | Florø | Norway | SCUBA | b | MUT | hotf | NR | standard | tele02 | 41 | 4.a |

**S1.2** Additional geographic information for samples sequenced where Ocean basins are as follows: 'NAt' = North Atlantic, 'NPa' = North Pacific, and 'Ind' = Indian.

| Dive Site Name | Location | Country | Ocean Basin | ICES Ecoregion | ICES Area |
|---|---|---|---|---|---|
| Mount Batten, Plymouth | Plymouth | England | NAt | Celtic Seas | 7.e |
| South Beach 1, Studland Bay, Dorset | Dorset | England | NAt | Greater North Sea | 7.d |
| South Beach 2, Studland Bay, Dorset | Dorset | England | NAt | Greater North Sea | 7.d |
| SMS Bayern, Scapa Flow, Orkney | Orkney | Scotland | NAt | Celtic Seas | 4.a |
| SMS Brummer, Scapa Flow, Orkney | Orkney | Scotland | NAt | Celtic Seas | 4.a |
| Dukes Dock, Liverpool | Liverpool | England | NAt | Celtic Seas | 7.a |
| Wuddy Rocks, St. Abbs, Scotland | St. Abbs Head | Scotland | NAt | Greater North Sea | 4.b |
| White Heugh, St. Abbs, Scotland | St. Abbs Head | Scotland | NAt | Greater North Sea | 4.b |
| Bodega de choco | Boa Vista | Cape Verde | NAt | NA | NA |
| San Carlos Beach Wall, CA, USA | California | USA | NPa | NA | NA |
| South Beach, Gulf of Aqaba, Jordan | Gulf of Aqaba | Jordan | Ind | NA | NA |
| SL9F, Sodwana Bay, South Africa | Sodwana Bay | South Africa | Ind | NA | NA |
| SL9M, Sodwana Bay, South Africa | Sodwana Bay | South Africa | Ind | NA | NA |
| SL7F1, Sodwana Bay, South Africa | Sodwana Bay | South Africa | Ind | NA | NA |
| SL7F2, Sodwana Bay, South Africa | Sodwana Bay | South Africa | Ind | NA | NA |
| Radbod, Ørsta, Norway | Ørsta | Norway | NAt | Norwegian Sea | 2.a.2 |
| Måløy, Norway | Måløy | Norway | NAt | Greater North Sea | 4.a |
| Dogfish Walk, Bergen, Norway | Bergen | Norway | NAt | Greater North Sea | 4.a |
| Welheim, Florø, Norway | Florø | Norway | NAt | Greater North Sea | 4.a |

**Abbreviations used in Supplementary tables S2-S6.**

Supplementary tables 2-6 show calculations of MOTUs and taxa from each sequencing library analysed in this dataset.
Tables X.1 show the total MOTUs and total reads in the sequencing library; the total MOTUs with taxonomy assigned at ≥70% (**motus70**), the total reads with taxonomy assigned at ≥70% (**reads70**) as well as the percent of each relative to the whole sequencing run (**percentmotus70**, **percentreads70**); the exact same statistics are calculated again but for MOTUs and reads with taxonomy assigned at ≥98% (i.e., **motus98**, **reads98**, **percentmotus98**, **percentreads98**).

Tables X.2 present statistics showing how MOTUs with taxonomy ≥70% were assigned. Specifically, the number of MOTUs with species-level assignments that had consensus between the three taxonomic assignment approaches (See Materials and Methods) (**species_3way**) and the percent of the total sequencing run that they represented (**percentmotuspecies_3way**). The same statistics were calculated at genus-level (**genus_3way**, **percentmotugenus_3way**). The number of MOTUs with species-level assignments that had consensus between two taxonomic assignment approaches (**species_2way**) and the percent of the total sequencing run that they represented (**percentmotuspecies_2way**). The number of MOTUs assigned to *Homo sapiens* (**human**) and the percent of the total sequencing run that they represented (**percentmotu_human**). And finally, the number of MOTUs with species-level assignments that did not have consensus were assigned using ecotag and a global reference database (**ecotag**) and the percent of the total sequencing run that they represented (**percentmotu_ecotag**).

Tables X.3 present statistics showing MOTUs with taxonomy ≥98% (**motus98**) at species (**species_98**) genus (**genus_98**) and higher (**other_98**) levels, the percent of the run they represent (**percentmotuspecies_98**, **percentmotugenus_98**, **percentmotuother_98**), and how they were assigned. Specifically, the number of MOTUs with genus and species-level assignments that had consensus between the three taxonomic assignment approaches (See Materials and Methods) (**method3way98**) and the percent of the total sequencing run that they represented (**percentmotu_3way98**). The number of MOTUs with genus and species-level assignments that had consensus between two taxonomic assignment approaches (**method2way98**) and the percent of the total sequencing run that they represented (**percentmotu_2way98**). The number of MOTUs assigned to *Homo sapiens* (**human98**) and the percent of the total sequencing run that they represented (**percentmotu_human98**). And finally, the number of MOTUs with species-level assignments that did not have consensus were assigned using ecotag and a global reference database (**ecotag98**) and the percent of the total sequencing run that they represented (**percentmotu_ecotag98**).

**Table S2.**

Sequencing statistics from sequencing run 1, library 1 (tele02 primer).

**S2.1.** Sequencing library reads and MOTUs after filtering by sequences which could be assigned taxonomy at 70% percent identity or 98% percent identity or higher.

| stat_all | value_all | stat_70 | value_70 | stat_98 | value_98 |
|---|---|---|---|---|---|
| total_MOTUs | 483 | motus70 | 451 | motus98 | 243 |
| Library_Reads | **1824604** | reads70 | 1823704 | reads98 | **1687590** |
| - | - | percentmotus70 | 93.37 | percentmotus98 | 50.31 |
| - | - | percentreads70 | 99.95 | percentreads98 | **92.49** |

**S2.2**

| stat | value |
|---|---|
| motus70 | 451 |
| species_3way | 64 |
| percentmotuspecies_3way | 14.19 |
| genus_3way | 7 |
| percentmotugenus_3way | 1.55 |
| species_2way | 153 |
| percentmotuspecies_2way | 33.92 |
| human | 31 |
| percentmotu_human | 6.87 |
| ecotag | 196 |
| percentmotu_ecotag | 43.46 |

**S2.3**

| stat2 | value2 |
|---|---|
| motus98 | 243 |
| species_98 | 204 |
| percentmotuspecies_98 | 83.95 |
| genus_98 | 27 |
| percentmotugenus_98 | 11.11 |
| other_98 | 12 |
| percentmotuother_98 | 4.94 |
| method3way98 | 71 |
| percentmotu_3way98 | **29.22** |
| method2way98 | 92 |
| percentmotu_2way98 | **37.86** |
| human98 | 6 |
| percentmotu_human98 | 2.47 |
| ecotag98 | 74 |
| percentmotu_ecotag98 | **30.45** |

**Table S3.**
Sequencing statistics from sequencing run 2, library 1 (elas02 primer).

**S3.1**

| stat_all | value_all | stat_70 | value_70 | stat_98 | value_98 |
|---|---|---|---|---|---|
| total_motus | 1324 | motus70 | 704 | motus98 | 450 |
| lib_reads | **954294** | reads70 | 927707 | reads98 | **884808** |
| - | - | percentmotus70 | 53.17 | percentmotus98 | 33.99 |
| - | - | percentreads70 | 97.21 | percentreads98 | **92.72** |

**S3.2**

| stat | value |
|---|---|
| motus70 | 704 |
| species_3way | 139 |
| percentmotuspecies_3way | 19.74 |
| genus_3way | 67 |
| percentmotugenus_3way | 9.52 |
| species_2way | 349 |
| percentmotuspecies_2way | 49.57 |
| human | 27 |
| percentmotu_human | 3.84 |
| ecotag | 122 |
| percentmotu_ecotag | 17.33 |

**S3.3**

| stat2 | value2 |
|---|---|
| motus98 | 450 |
| species_98 | 360 |
| percentmotuspecies_98 | 80 |
| genus_98 | 75 |
| percentmotugenus_98 | 16.67 |
| other_98 | 15 |
| percentmotuother_98 | 3.33 |
| method3way98 | 151 |
| percentmotu_3way98 | **33.56** |
| method2way98 | 242 |
| percentmotu_2way98 | **53.78** |
| human98 | 7 |
| percentmotu_human98 | 1.56 |
| ecotag98 | 50 |
| percentmotu_ecotag98 | **11.11** |

**Table S4.**

Sequencing statistics from sequencing run 2, library 2 (tele02 primer).

**S4.1**

| stat_all | value_all | stat_70 | value_70 | stat_98 | value_98 |
|---|---|---|---|---|---|
| total_motus | 828 | motus70 | 820 | motus98 | 385 |
| lib_reads | **1494951** | reads70 | 1494884 | reads98 | **1292245** |
| - | - | percentmotus70 | 99.03 | percentmotus98 | 46.50 |
| - | - | percentreads70 | 100.00 | percentreads98 | **86.44** |

**S4.2**

| stat | value |
|---|---|
| motus70 | 820 |
| species_3way | 121 |
| percentmotuspecies_3way | 14.76 |
| genus_3way | 69 |
| percentmotugenus_3way | 8.41 |
| species_2way | 177 |
| percentmotuspecies_2way | 21.59 |
| human | 62 |
| percentmotu_human | 7.56 |
| ecotag | 391 |
| percentmotu_ecotag | 47.68 |

**S4.3**

| stat2 | value2 |
|---|---|
| motus98 | 385 |
| species_98 | 252 |
| percentmotuspecies_98 | 65.45 |
| genus_98 | 114 |
| percentmotugenus_98 | 29.61 |
| other_98 | 19 |
| percentmotuother_98 | 4.94 |
| method3way98 | 126 |
| percentmotu_3way98 | **32.73** |
| method2way98 | 127 |
| percentmotu_2way98 | **32.99** |
| human98 | 9 |
| percentmotu_human98 | 2.34 |
| ecotag98 | 123 |
| percentmotu_ecotag98 | **31.95** |

**Table S5.**
Sequencing statistics from sequencing run 3, library 1 (elas02 primer).

**S5.1**

| stat_all | value_all | stat_70 | value_70 | stat_98 | value_98 |
|---|---|---|---|---|---|
| total_motus | 3784 | motus70 | 2493 | motus98 | 917 |
| lib_reads | **909806** | reads70 | 895852 | reads98 | **785300** |
| - | - | percentmotus70 | 65.88266 | percentmotus98 | 24.23362 |
| - | - | percentreads70 | 98.46627 | percentreads98 | **86.3151** |

**S5.2**

| stat | value |
|---|---|
| motus70 | 2493 |
| species_3way | 215 |
| percentmotuspecies_3way | 8.62 |
| genus_3way | 73 |
| percentmotugenus_3way | 2.93 |
| species_2way | 651 |
| percentmotuspecies_2way | 26.11 |
| human | 402 |
| percentmotu_human | 16.13 |
| ecotag | 1152 |
| percentmotu_ecotag | 46.21 |

**S5.3**

| stat2 | value2 |
|---|---|
| motus98 | 917 |
| species_98 | 716 |
| percentmotuspecies_98 | 78.08 |
| genus_98 | 147 |
| percentmotugenus_98 | 16.03 |
| other_98 | 54 |
| percentmotuother_98 | 5.89 |
| method3way98 | 213 |
| percentmotu_3way98 | **23.23** |
| method2way98 | 487 |
| percentmotu_2way98 | **53.11** |
| human98 | 48 |
| percentmotu_human98 | 5.23 |
| ecotag98 | 169 |
| percentmotu_ecotag98 | **18.43** |

**Table S6.**

Sequencing statistics from sequencing run 4, library 1 (tele02 primer).

**S6.1**

| stat_all | value_all | stat_70 | value_70 | stat_98 | value_98 |
|---|---|---|---|---|---|
| total_motus | 1071 | motus70 | 1064 | motus98 | 333 |
| lib_reads | **989266** | reads70 | 989113 | reads98 | **929358** |
| - | - | percentmotus70 | 99.35 | percentmotus98 | 31.09 |
| - | - | percentreads70 | 99.98 | percentreads98 | **93.94** |

**S6.2**

| stat | value |
|---|---|
| motus70 | 1064 |
| species_3way | 117 |
| percentmotuspecies_3way | 11.00 |
| genus_3way | 39 |
| percentmotugenus_3way | 3.67 |
| species_2way | 301 |
| percentmotuspecies_2way | 28.29 |
| human | 179 |
| percentmotu_human | 16.82 |
| ecotag | 428 |
| percentmotu_ecotag | 40.23 |

**S6.4**

| stat2 | value2 |
|---|---|
| motus98 | 333 |
| species_98 | 310 |
| percentmotuspecies_98 | 93.09 |
| genus_98 | 20 |
| percentmotugenus_98 | 6.01 |
| other_98 | 3 |
| percentmotuother_98 | 0.90 |
| method3way98 | 125 |
| percentmotu_3way98 | **37.54** |
| method2way98 | 175 |
| percentmotu_2way98 | **52.55** |
| human98 | 16 |
| percentmotu_human98 | 4.80 |
| ecotag98 | 17 |
| percentmotu_ecotag98 | **5.11** |

**Table S7.**
Taxonomic assignments and taxa category of teleosts detected in the Ocean Exhibit.

| Teleost Detection | Category |
|---|---|
| *Abudefduf* | inventory |
| *Acanthurus* | inventory |
| *Anisotremus virginicus* | inventory |
| *Balistes vetula* | inventory |
| *Caesio caerulaurea* | inventory |
| *Caesio cuning* | inventory |
| *Chrysiptera cyanea* | inventory |
| *Coris gaimard* | inventory |
| *Diodon* | inventory |
| *Epinephelus* | inventory |
| *Epinephelus lanceolatus* | inventory |
| *Gnathanodon speciosus* | inventory |
| *Grammistes sexlineatus* | inventory |
| *Gymnothorax funebris* | inventory |
| *Haemulon sciurus* | inventory |
| *Heniochus acuminatus* | inventory |
| *Labroides dimidiatus* | inventory |
| *Lutjanus* | inventory |
| *Lutjanus kasmira* | inventory |
| *Lutjanus sebae* | inventory |
| *Megalops atlanticus* | inventory |
| *Megalops cyprinoides* | inventory |
| *Naso* | inventory |
| *Odonus niger* | inventory |
| *Paracanthurus hepatus* | inventory |
| *Paranthias colonus* | inventory |
| *Platax* | inventory |
| *Pomacanthus maculosus* | inventory |
| *Pseudobalistes fuscus* | inventory |
| *Siganus* | inventory |
| *Thalassoma* | inventory |
| *Argyrosomus regius* | putative inventory |
| *Monodactylus argenteus* | putative inventory |
| *Naso elegans* | putative inventory |
| *Pseudocaranx dentex* | putative inventory |
| *Trachinotus blochii* | putative inventory |
| *Trachinotus rhodopus* | putative inventory |
| *Ammodytes* | food |
| *Auxis* | food |

| | |
|---|---|
| *Clupea harengus* | food |
| *Clupeinae* | food |
| *Euthynnus* | food |
| *Gadidae* | food |
| *Gadus* | food |
| *Hippoglossus* | food |
| *Melanogrammus* | food |
| *Merluccius merluccius* | food |
| *Sardina pilchardus* | food |
| *Scomber* | food |
| *Scomber scombrus* | food |
| *Trachurus trachurus* | food |

**Table S8.**

Taxonomic assignments and taxa category of teleosts detected in the Coral cave exhibit.

| Teleost Detection | Category |
|---|---|
| Abudefduf | inventory |
| Acanthurus | inventory |
| Acanthurus olivaceus | inventory |
| Arothron | inventory |
| Arothron hispidus | inventory |
| Balistoides conspicillum | inventory |
| Chaetodon | inventory |
| Chrysiptera | inventory |
| Chrysiptera cyanea | inventory |
| Diodon | inventory |
| Gymnomuraena zebra | inventory |
| Gymnothorax flavimarginatus | inventory |
| Holacanthus ciliaris | inventory |
| Labroides dimidiatus | inventory |
| Melichthys | inventory |
| Myripristis | inventory |
| Odonus niger | inventory |
| Paracanthurus hepatus | inventory |
| Parupeneus cyclostomus | inventory |
| Platax | inventory |
| Plectorhinchus vittatus | inventory |
| Pomacanthus | inventory |
| Pseudanthias squamipinnis | inventory |
| Siganus | inventory |
| Thalassoma | inventory |
| Zebrasoma | inventory |
| Zebrasoma desjardinii | inventory |
| Balistapus undulatus | putative inventory |
| Cephalopholis sexmaculata | putative inventory |
| Chaetodon auriga | putative inventory |
| Chromileptes altivelis | putative inventory |
| Diagramma | putative inventory |
| Gymnothorax kidako | putative inventory |
| Hemitaurichthys | putative inventory |
| Naso elegans | putative inventory |
| Zebrasoma velifer | putative inventory |
| Clupea harengus | food |
| Gadidae | food |
| Micromesistius poutassou | food |
| Pollachius virens | food |

| | |
|---|---|
| Scomber | food |
| Scomberesox saurus | food |

**Table S9.**

Results of the linear models run on the timed soaking experiments and controlled aquarium dives.

**S9.1** Linear model results for change in MOTUs through time for the soaking experiment.

| Coefficients | | | | |
|---|---|---|---|---|
| | estimate | Std. error | t value | Pr(>\|t\|) |
| Intercept (MOTUs) | 16.68991 | 4.67073 | 3.573 | 0.00217 |
| Slope (time) | 0.12402 | 0.03774 | 3.287 | 0.00410 |
| | Multiple $R^2$: 0.375 | Adjusted $R^2$: 0.3403 | F statistic : 10.8 on 1 and 18 | p-value: **0.004101** |

**S9.2** Linear model results for change in taxa through time for the soaking experiment.

| Coefficients | | | | |
|---|---|---|---|---|
| | estimate | Std. error | t value | Pr(>\|t\|) |
| Intercept (Taxa) | 10.28326 | 2.09996 | 4.897 | 0.000116 |
| Slope (time) | 0.08877 | 0.01697 | 5.232 | 5.63e-05 |
| | Multiple $R^2$: 0.6033 | Adjusted $R^2$: 0.5813 | F statistic : 27.38 on 1 and 18 | p-value: **5.633e-05** |

**S9.3** Linear model results for change in MOTUs through time for the soaking experiment and with Ocean Exhibit dives added in.

| Coefficients | | | | |
|---|---|---|---|---|
| | estimate | Std. error | t value | Pr(>\|t\|) |
| Intercept (Taxa) | 23.28414 | 4.47017 | 5.209 | 1.94e-05 |
| Slope (time) | 0.09958 | 0.04097 | 2.431 | 0.0223 |
| | Multiple $R^2$: 0.1851 | Adjusted $R^2$: 0.1538 | F statistic : 5.908 on 1 and 26 DF | p-value: **0.02228** |

**S9.3** Linear model results for change in taxa through time for the soaking experiment and with Ocean Exhibit dives added in.

| Coefficients | | | | |
|---|---|---|---|---|
| | estimate | Std. error | t value | Pr(>\|t\|) |
| Intercept (Taxa) | 14.78161 | 2.40377 | 6.149 | 1.68e-06 |
| Slope (time) | 0.07179 | 0.02203 | 3.258 | 0.00312 |
| | Multiple $R^2$: 0.2899 | Adjusted $R^2$: 0.2626 | F statistic : 10.62 on 1 and 26 DF | p-value: **0.003116** |

**Table S10.**

Results of PERMANOVA and beta-dispersion tests comparing syringe-filter eDNA samples to diver metaprobes (relating to Fig. 4E).

**S10.1** Permutation test (function: permutest) for homogeneity of multivariate dispersions for dive sites.

| | DF | Sum Sq | Mean Sq | F | N.perm | Pr(>F) |
|---|---|---|---|---|---|---|
| Groups | 3 | 0.65627 | 0.218755 | 9.2787 | 999 | 0.001 |
| Residuals | 32 | 0.75443 | 0.023576 | | | |

**S10.2** Permutation test (function: permutest) for homogeneity of multivariate dispersions for sample type.

| | DF | Sum Sq | Mean Sq | F | N.perm | Pr(>F) |
|---|---|---|---|---|---|---|
| Groups | 1 | 0.02022 | 0.020219 | 0.8 | 999 | 0.386 |
| Residuals | 34 | 0.85925 | 0.025272 | | | |

**S10.3** PERMANOVA (function: adonis) testing. adonis2(jac_dat ~ site.name, data = cem_jac, permutations = 999, strata = cem_jac$type)

| | DF | SumOfSqs | R2 | F | Pr(>F) |
|---|---|---|---|---|---|
| type | 3 | 3.9786 | 0.3041 | 4.6611 | 0.001 |
| Residual | 32 | 9.1047 | 0.6959 | | |
| total | 35 | 13.0833 | 1.0000 | | |

**S10.4** PERMANOVA (function: adonis) testing. adonis2(jac_dat ~ type, data = cem_jac, permutations = 999, strata = cem_jac$site.name)

| | DF | SumOfSqs | R2 | F | Pr(>F) |
|---|---|---|---|---|---|
| type | 1 | 0.5922 | 0.04526 | 1.6118 | 0.057 |
| Residual | 34 | 12.4911 | 0.95474 | | |
| total | 35 | 13.0833 | 1.00000 | | |

**Table S11.**

GLMM output testing if input weight of gauze and lysis buffer effects alpha-diversity.

| | | | | |
|---|---|---|---|---|
| REML criterion at convergence | 109.9 | | | |
| Scaled residuauls: | | | | |
| Min | 1Q | Median | 3Q | Max |
| -1.4190 | -0.4807 | -0.2409 | 0.3736 | 2.6272 |
| Random effects: | | | | |
| Groups | Name | Variance | Standard deviation | Corr |
| mp_id | (intercept) | 2.321e+01 | 4.8177739 | |
| | Weight_g | 2.322e+01 | 4.8184723 | 0.00 |
| | lysis | 4.791e-07 | 0.0006922 | -0.01 -1.00 |
| | Residual | 2.322e+01 | 4.8185362 | |
| Number of obs: | 18 | Groups: | mp_id, 2 | |
| Fixed effects: | | | | |

| | Estimate | Std. Error | df | T value | Pr(>\|t\|) |
|---|---|---|---|---|---|
| (intercept) | -16.332898 | 48.364102 | 2.030992 | -0.338 | 0.767 |
| Weight_g | 3.878243 | 8.048875 | 1.173324 | 0.482 | 0.704 |
| lysis | -0.001883 | 0.001932 | 4.284934 | -0.975 | 0.382 |
| time | 0.255225 | 0.577383 | 2.015853 | 0.442 | 0.701 |

| Correlation of fixed effects: | | | | |
|---|---|---|---|---|
| | | intercept | Weight_g | lysis |
| | Weight_g | -0.016 | | |
| | lysis | 0.003 | -0.876 | |
| | time | -0.996 | -0.014 | 0.007 |

**Table S12.**

GLMM to test for differences in extraction method.

| | | | | | |
|---|---|---|---|---|---|
| REML criterion at convergence | | 68.6 | | | |
| Scaled residuauls: | | | | | |
| Min | | 1Q | Median | 3Q | Max |
| -1.68974 | | -0.70852 | -0.06431 | 0.52593 | 2.90766 |
| Random effects: | | | | | |
| Groups | | Name | Variance | Standard deviation | Corr |
| mp_id | | (intercept) | 2.062e-10 | 1.436e-05 | |
| | | ExtractionQBT | 6.721e-10 | 2.592e-05 | -0.91 |
| | | Residual | 3.692e-01 | 6.076e-01 | |
| Number of obs: | | 36 | Groups: | mp_id, 7 | |
| Fixed effects: | | | | | |
| | Estimate | Std. Error | df | T value | Pr(>\|t\|) |
| (intercept) | 1.02673 | 0.20254 | 32 | 5.069 | 1.62e-05 |
| extractionQBT | 0.08671 | 0.20254 | 32 | 0.428 | 0.671 |
| Site.names Bayern, Orkney | -0.15793 | 0.24806 | 32 | -0.637 | 0.529 |
| Site.names Beach 1, Studland Bay | -0.36166 | 0.24806 | 32 | -1.458 | 0.155 |
| Correlation of fixed effects: | | | | | |
| | | | intercept | extractionQBT | Site.names Bayern, Orkney |
| | | extractionQBT | -0.500 | | |
| | | Site.names Bayern, Orkney | -0.612 | 0.000 | |
| | | Site.names Beach 1, Studland Bay | -0.612 | 0.000 | 0.500 |

**Table S13.**

GLMM to test differences in preservation method.

| | | | | |
|---|---|---|---|---|
| REML criterion at convergence | 138 | | | |
| Scaled residuauls: | | | | |
| Min | 1Q | Median | 3Q | Max |
| -1.23162 | -0.70790 | -0.09452 | 0.61072 | 1.92178 |
| Random effects: | | | | |
| Groups | Name | Variance | Standard deviation | Corr |
| mp_id | (intercept) | 0.0000 | 0.0000 | |
| | preservationEt | 0.7409 | 0.8608 | NaN |
| | Residual | 7.2981 | 2.7015 | |
| Number of obs: | 33 | Groups: | mp_id, 15 | |
| Fixed effects: | | | | |

| | Estimate | Std. Error | df | T value | Pr(>\|t\|) |
|---|---|---|---|---|---|
| (intercept) | -1.4699 | 6.2688 | 12.9708 | -0.234 | 0.81827 |
| preservationEt | 3.0235 | 0.9993 | 25.3678 | 3.026 | **0.00562** |
| Site.names Bayern, Orkney | 3.3037 | 3.3489 | 17.9087 | 0.987 | 0.33702 |
| Site.names Brummer, Orkney | 0.9473 | 1.6823 | 25.2057 | 0.563 | 0.57836 |
| Site.names White Heugh, St.Abbs | -0.1288 | 1.5969 | 24.6572 | -0.081 | 0.93636 |
| Site.names Wuddy Rocks, St.Abbs | -0.9541 | 1.5969 | 24.6572 | -0.597 | 0.55565 |
| Sequence.run | 1.8409 | 3.0290 | 13.0953 | 0.608 | 0.55375 |

Correlation of fixed effects:

| | intercept | preservationEt | Site.names Bayern, Orkney | Site.names Brummer, Orkney | Site.names White Heugh, St.Abbs | Site.names Wuddy Rocks, St.Abbs |
|---|---|---|---|---|---|---|
| preservationEt | -0.260 | | | | | |
| Site.names Bayern, Orkney | -0.936 | 0.128 | | | | |
| Site.names Brummer, Orkney | 0.188 | -0.060 | -0.056 | | | |
| Site.names White Heugh, St.Abbs | -0.127 | 0.000 | 0.238 | 0.475 | | |
| Site.names Wuddy Rocks, St.Abbs | -0.127 | 0.000 | 0.238 | 0.475 | 0.500 | |
| Sequence.run | -0.981 | 0.190 | 0.896 | -0.315 | 0.000 | 0.000 |

**Table S14.**

Results of CCA for diver metaprobe data.

**S14.1** Metadata variables included for CCA model building process of all diver metaprobe data.

| Metadata variable | Description |
|---|---|
| latitude | Continuous, decimal degrees |
| preservation | Categorical, indicating silica beads or ethanol |
| extraction | Categorical, indicating DNA extraction method |
| Weight_class | Categorical, indicating input weight range |
| Primer | Categorical, indicating primer used for PCR |
| Sequence.run | Categorical, indicating sequence run sample is from |
| Ocean_basin | Categorical, indicating ocean basin |

**S14.2** Significance of the variables selected for the minimum adequate CCA model of all data.

| | DF | Chi Square | F | Pr(>F) |
|---|---|---|---|---|
| Ocean_basin | 2 | 1.9760 | 5.0265 | **0.001** |
| latitude | 1 | 0.8184 | 4.1638 | **0.001** |
| residual | 108 | 21.2283 | | |

**S14.3** Metadata variables included for model building process of North Atlantic data.

| Metadata variable | Description |
|---|---|
| latitude | Continuous, decimal degrees |
| preservation | Categorical, indicating silica beads or ethanol |
| extraction | Categorical, indicating DNA extraction method |
| Weight_class | Categorical, indicating input weight range |
| Primer | Categorical, indicating primer used for PCR |
| Sequence.run | Categorical, indicating sequence run sample is from |
| Ices_ecor | Categorical, ICES ecoregion |
| Ices_area | Categorical, ICES area |

**S14.4** Significance of the variables selected for the minimum adequate CCA model of North Atlantic data.

| | DF | Chi Square | F | Pr(>F) |
|---|---|---|---|---|
| Ices_area | 5 | 3.0978 | 5.1515 | **0.001** |
| latitude | 1 | 0.1656 | 1.3772 | 0.190 |
| residual | 85 | 10.2229 | | |

**Table S15.**

Species-level detections from metaprobes worn by divers in nature. IUCN categories for each species are given, where 'CR' = critically endangered, 'EN' = endangered, 'VU' = vulnerable, 'NT' = near threatened, 'DD' = data deficient, 'LC' = least concern, and 'NE' = not evaluated.

| Number | Class | Species | Read Counts | IUCN |
|---|---|---|---|---|
| 1 | Actinopterygii | *Anguilla anguilla* | 2360 | **CR** |
| 2 | Actinopterygii | *Gadus morhua* | 1141 | **VU** |
| 3 | Actinopterygii | *Mola mola* | 198 | **VU** |
| 4 | Actinopterygii | *Trachurus trachurus* | 93 | **VU** |
| 5 | Actinopterygii | *Bodianus pulcher* | 46 | **VU** |
| 6 | Actinopterygii | *Sardinella maderensis* | 4 | **VU** |
| 7 | Actinopterygii | *Salmo salar* | 55188 | **NT** |
| 8 | Actinopterygii | *Pagellus bogaraveo* | 259 | **NT** |
| 9 | Actinopterygii | *Solea solea* | 548 | **DD** |
| 10 | Actinopterygii | *Engraulis mordax* | 267 | **DD** |
| 11 | Actinopterygii | *Brachyistius frenatus* | 43 | **DD** |
| 12 | Actinopterygii | *Ammodytes tobianus* | 39 | **DD** |
| 13 | Actinopterygii | *Dicentrarchus labrax* | 123089 | LC |
| 14 | Actinopterygii | *Trisopterus minutus* | 122708 | LC |
| 15 | Actinopterygii | *Atherina boyeri* | 81556 | LC |
| 16 | Actinopterygii | *Trisopterus esmarkii* | 80690 | LC |
| 17 | Actinopterygii | *Gasterosteus aculeatus* | 47524 | LC |
| 18 | Actinopterygii | *Symphodus melops* | 25903 | LC |
| 19 | Actinopterygii | *Labrus bergylta* | 19641 | LC |
| 20 | Actinopterygii | *Symphodus bailloni* | 16254 | LC |
| 21 | Actinopterygii | *Pomatoschistus minutus* | 15600 | LC |
| 22 | Actinopterygii | *Sprattus sprattus* | 13969 | LC |
| 23 | Actinopterygii | *Chelon auratus* | 13652 | LC |
| 24 | Actinopterygii | *Pseudanthias squamipinnis* | 12871 | LC |
| 25 | Actinopterygii | *Taurulus bubalis* | 12243 | LC |
| 26 | Actinopterygii | *Atherinopsis californiensis* | 10133 | LC |
| 27 | Actinopterygii | *Ctenolabrus rupestris* | 9435 | LC |
| 28 | Actinopterygii | *Sardinops sagax* | 4985 | LC |
| 29 | Actinopterygii | *Pollachius pollachius* | 4073 | LC |
| 30 | Actinopterygii | *Porichthys notatus* | 3534 | LC |
| 31 | Actinopterygii | *Gobiusculus flavescens* | 2083 | LC |
| 32 | Actinopterygii | *Gobius paganellus* | 1942 | LC |
| 33 | Actinopterygii | *Gymnammodytes semisquamatus* | 1729 | LC |
| 34 | Actinopterygii | *Clupea harengus* | 1716 | LC |
| 35 | Actinopterygii | *Gobius niger* | 1580 | LC |
| 36 | Actinopterygii | *Platichthys stellatus* | 1561 | LC |

| | | | | |
|---|---|---|---|---|
| 37 | Actinopterygii | *Syngnathus typhle* | 1486 | LC |
| 38 | Actinopterygii | *Oxylebius pictus* | 1455 | LC |
| 39 | Actinopterygii | *Lutjanus bohar* | 1264 | LC |
| 40 | Actinopterygii | *Naso lopezi* | 1154 | LC |
| 41 | Actinopterygii | *Scarus rubroviolaceus* | 966 | LC |
| 42 | Actinopterygii | *Nerophis ophidion* | 822 | LC |
| 43 | Actinopterygii | *Syngnathus acus* | 785 | LC |
| 44 | Actinopterygii | *Acanthurus nigrofuscus* | 767 | LC |
| 45 | Actinopterygii | *Parablennius gattorugine* | 642 | LC |
| 46 | Actinopterygii | *Coryphoblennius galerita* | 612 | LC |
| 47 | Actinopterygii | *Ciliata mustela* | 587 | LC |
| 48 | Actinopterygii | *Spinachia spinachia* | 557 | LC |
| 49 | Actinopterygii | *Scophthalmus rhombus* | 509 | LC |
| 50 | Actinopterygii | *Sufflamen chrysopterum* | 499 | LC |
| 51 | Actinopterygii | *Sparus aurata* | 452 | LC |
| 52 | Actinopterygii | *Pterocaesio marri* | 427 | LC |
| 53 | Actinopterygii | *Balistapus undulatus* | 420 | LC |
| 54 | Actinopterygii | *Pomatoschistus pictus* | 393 | LC |
| 55 | Actinopterygii | *Phanerodon vacca* | 388 | LC |
| 56 | Actinopterygii | *Labrus mixtus* | 380 | LC |
| 57 | Actinopterygii | *Pomatoschistus microps* | 339 | LC |
| 58 | Actinopterygii | *Conger conger* | 292 | LC |
| 59 | Actinopterygii | *Pervagor janthinosoma* | 289 | LC |
| 60 | Actinopterygii | *Lipophrys pholis* | 288 | LC |
| 61 | Actinopterygii | *Acanthurus leucosternon* | 271 | LC |
| 62 | Actinopterygii | *Paracaesio sordida* | 271 | LC |
| 63 | Actinopterygii | *Cirrhitichthys oxycephalus* | 267 | LC |
| 64 | Actinopterygii | *Phoxinus phoxinus* | 253 | LC |
| 65 | Actinopterygii | *Merluccius merluccius* | 252 | LC |
| 66 | Actinopterygii | *Pycnochromis nigrurus* | 241 | LC |
| 67 | Actinopterygii | *Embiotoca jacksoni* | 226 | LC |
| 68 | Actinopterygii | *Acanthurus thompsoni* | 203 | LC |
| 69 | Actinopterygii | *Syngnathus rostellatus* | 200 | LC |
| 70 | Actinopterygii | *Anampses caeruleopunctatus* | 168 | LC |
| 71 | Actinopterygii | *Bodianus bilunulatus* | 142 | LC |
| 72 | Actinopterygii | *Callionymus lyra* | 125 | LC |
| 73 | Actinopterygii | *Sardina pilchardus* | 125 | LC |
| 74 | Actinopterygii | *Rhacochilus vacca* | 122 | LC |
| 75 | Actinopterygii | *Myripristis kuntee* | 114 | LC |
| 76 | Actinopterygii | *Sillago sihama* | 111 | LC |
| 77 | Actinopterygii | *Ostorhinchus apogonoides* | 109 | LC |
| 78 | Actinopterygii | *Anarrhichthys ocellatus* | 95 | LC |
| 79 | Actinopterygii | *Odonus niger* | 83 | LC |

| | | | | |
|---|---|---|---|---|
| 80 | Actinopterygii | *Gymnothorax nudivomer* | 79 | LC |
| 81 | Actinopterygii | *Chlorophthalmus agassizi* | 72 | LC |
| 82 | Actinopterygii | *Citharichthys stigmaeus* | 67 | LC |
| 83 | Actinopterygii | *Labroides dimidiatus* | 66 | LC |
| 84 | Actinopterygii | *Coris caudimacula* | 60 | LC |
| 85 | Actinopterygii | *Diplodus sargus* | 52 | LC |
| 86 | Actinopterygii | *Lutjanus fulviflamma* | 47 | LC |
| 87 | Actinopterygii | *Liparis montagui* | 46 | LC |
| 88 | Actinopterygii | *Cyclothone alba* | 44 | LC |
| 89 | Actinopterygii | *Belone belone* | 42 | LC |
| 90 | Actinopterygii | *Cantherhines dumerilii* | 41 | LC |
| 91 | Actinopterygii | *Girella nigricans* | 40 | LC |
| 92 | Actinopterygii | *Zoarces viviparus* | 38 | LC |
| 93 | Actinopterygii | *Siganus luridus* | 35 | LC |
| 94 | Actinopterygii | *Parupeneus macronemus* | 33 | LC |
| 95 | Actinopterygii | *Ciliata septentrionalis* | 31 | LC |
| 96 | Actinopterygii | *Chromis weberi* | 22 | LC |
| 97 | Actinopterygii | *Scomber scombrus* | 22 | LC |
| 98 | Actinopterygii | *Acanthurus dussumieri* | 21 | LC |
| 99 | Actinopterygii | *Halichoeres californicus* | 20 | LC |
| 100 | Actinopterygii | *Halichoeres cosmetus* | 18 | LC |
| 101 | Actinopterygii | *Gobio gobio* | 17 | LC |
| 102 | Actinopterygii | *Gymnothorax melatremus* | 17 | LC |
| 103 | Actinopterygii | *Sparisoma cretense* | 17 | LC |
| 104 | Actinopterygii | *Cephalopholis sexmaculata* | 16 | LC |
| 105 | Actinopterygii | *Azurina lepidolepis* | 15 | LC |
| 106 | Actinopterygii | *Micrenophrys lilljeborgii* | 15 | LC |
| 107 | Actinopterygii | *Citharichthys sordidus* | 14 | LC |
| 108 | Actinopterygii | *Lutjanus argentimaculatus* | 13 | LC |
| 109 | Actinopterygii | *Pomacanthus imperator* | 12 | LC |
| 110 | Actinopterygii | *Salmo trutta* | 12 | LC |
| 111 | Actinopterygii | *Cymatogaster aggregata* | 11 | LC |
| 112 | Actinopterygii | *Heterostichus rostratus* | 11 | LC |
| 113 | Actinopterygii | *Caranx crysos* | 10 | LC |
| 114 | Actinopterygii | *Centropyge multispinis* | 10 | LC |
| 115 | Actinopterygii | *Crossorhombus valderostratus* | 10 | LC |
| 116 | Actinopterygii | *Lutjanus kasmira* | 10 | LC |
| 117 | Actinopterygii | *Parupeneus pleurostigma* | 10 | LC |
| 118 | Actinopterygii | *Megalaspis cordyla* | 9 | LC |
| 119 | Actinopterygii | *Caesio varilineata* | 8 | LC |
| 120 | Actinopterygii | *Ptereleotris evides* | 7 | LC |
| 121 | Actinopterygii | *Plectorhinchus orientalis* | 6 | LC |
| 122 | Actinopterygii | *Sargocentron diadema* | 6 | LC |

| | | | | |
|---|---|---|---|---|
| 123 | Actinopterygii | *Antennatus tuberosus* | 5 | LC |
| 124 | Actinopterygii | *Stethojulis interrupta* | 5 | LC |
| 125 | Actinopterygii | *Ctenochaetus binotatus* | 4 | LC |
| 126 | Actinopterygii | *Trachinocephalus myops* | 4 | LC |
| 127 | Actinopterygii | *Trachinotus blochii* | 4 | LC |
| 128 | Actinopterygii | *Gnathanodon speciosus* | 3 | LC |
| 129 | Actinopterygii | *Sufflamen fraenatum* | 3 | LC |
| 130 | Actinopterygii | *Pollachius virens* | 107912 | NE |
| 131 | Actinopterygii | *Ammodytes marinus* | 7591 | NE |
| 132 | Actinopterygii | *Pholis gunnellus* | 3025 | NE |
| 133 | Actinopterygii | *Chirolophis japonicus* | 2470 | NE |
| 134 | Actinopterygii | *Trisopterus luscus* | 1169 | NE |
| 135 | Actinopterygii | *Toxabramis swinhonis* | 1031 | NE |
| 136 | Actinopterygii | *Cyclopterus lumpus* | 812 | NE |
| 137 | Actinopterygii | *Molva molva* | 341 | NE |
| 138 | Actinopterygii | *Rhacochilus toxotes* | 217 | NE |
| 139 | Actinopterygii | *Gadiculus argenteus* | 184 | NE |
| 140 | Actinopterygii | *Oplegnathus fasciatus* | 158 | NE |
| 141 | Actinopterygii | *Scorpaenichthys marmoratus* | 85 | NE |
| 142 | Actinopterygii | *Liparis mucosus* | 64 | NE |
| 143 | Actinopterygii | *Echiichthys vipera* | 58 | NE |
| 144 | Actinopterygii | *Plectorhinchus flavomaculatus* | 55 | NE |
| 145 | Actinopterygii | *Esselenichthys carli* | 44 | NE |
| 146 | Actinopterygii | *Sebastes gilli* | 40 | NE |
| 147 | Actinopterygii | *Micrometrus aurora* | 35 | NE |
| 148 | Actinopterygii | *Myoxocephalus scorpius* | 34 | NE |
| 149 | Actinopterygii | *Ophiodon elongatus* | 34 | NE |
| 150 | Actinopterygii | *Sphyraena jello* | 30 | NE |
| 151 | Actinopterygii | *Oncorhynchus mykiss* | 28 | NE |
| 152 | Actinopterygii | *Micromesistius poutassou* | 13 | NE |
| 153 | Actinopterygii | *Anoplopoma fimbria* | 12 | NE |
| 154 | Actinopterygii | *Plotosus lineatus* | 6 | NE |
| 155 | Actinopterygii | *Clinocottus recalvus* | 3 | NE |
| 156 | Actinopterygii | *Micrometrus minimus* | 3 | NE |
| 157 | Aves | *Alca torda* | 4729 | LC |
| 158 | Aves | *Columba livia* | 1302 | LC |
| 159 | Aves | *Pelecanus occidentalis* | 125 | LC |
| 160 | Aves | *Arenaria interpres* | 101 | LC |
| 161 | Aves | *Columba palumbus* | 84 | LC |
| 162 | Aves | *Uria aalge* | 63 | LC |
| 163 | Aves | *Larus occidentalis* | 56 | LC |
| 164 | Elasmobranchii | *Rhynchobatus djiddensis* | 3 | **CR** |
| 165 | Elasmobranchii | *Himantura uarnak* | 1309 | **EN** |

| | | | | |
|---|---|---|---|---|
| 166 | Elasmobranchii | *Taeniurops meyeni* | 14158 | **VU** |
| 167 | Elasmobranchii | *Pateobatis fai* | 109 | **VU** |
| 168 | Elasmobranchii | *Pateobatis jenkinsii* | 13 | **VU** |
| 169 | Elasmobranchii | *Alopias vulpinus* | 9 | **VU** |
| 170 | Elasmobranchii | *Triaenodon obesus* | 6 | **VU** |
| 171 | Elasmobranchii | *Squalus acanthias* | 3 | **VU** |
| 172 | Elasmobranchii | *Myliobatis californica* | 15 | LC |
| 173 | Elasmobranchii | *Lamna ditropis* | 5 | LC |
| 174 | Mammalia | *Phocoena phocoena* | 227 | **VU** |
| 175 | Mammalia | *Tursiops aduncus* | 31 | **NT** |
| 176 | Mammalia | *Zalophus californianus* | 17933 | LC |
| 177 | Mammalia | *Mus musculus* | 1101 | LC |
| 178 | Mammalia | *Phoca vitulina* | 81 | LC |
| 179 | Mammalia | *Stenella attenuata* | 11 | LC |
| 180 | Reptilia | *Caretta caretta* | 5 | **VU** |
